# Foliar Application of Polymer-coated Manganese Dioxide Nanoparticles: Mechanisms of Uptake and Metabolic Responses in Manganese Deficient Barley

**DOI:** 10.1101/2025.08.11.669670

**Authors:** Andrea Pinna, Asbjørn Krarup Grønbæk, Emil Visby Kristensen, Noémie Thiébaut, Augusta Szameitat, Birte Martin-Bertelsen, Tue Hassenkam, Jean-Claude Grivel, Rajmund Mokso, Søren Husted

**Author notes:** Corresponding author: Søren Husted.

## Abstract

The application of nanotechnology in plant science is unlocking innovative approaches to enhance nutrient use efficiency in crops, particularly through foliar fertilization. This study demonstrates that colloidally stable, pH-responsive polyacrylic acid (PAA)-coated manganese dioxide (MnO_2_) nanoparticles (nPAA-MnO_2_) can be designed to significantly restore key metabolic functionalities in manganese (Mn)-deficient barley (*Hordeum vulgare*) within a few days. Using a combination of advanced bioimaging techniques - including confocal laser scanning microscopy (CLSM), laser ablation-inductively coupled plasma-mass spectrometry (LA-ICP-MS), and X-ray nano-computed tomography (nano-CT), we mapped the uptake and distribution pathways of nPAA-MnO_2_ compared to ionic Mn. While soluble Mn^2+^ ions primarily enter through hydrophilic cuticular pores, nPAA-MnO_2_ penetrates leaves *via* stomata, facilitated by the application of an organosilicone surfactant and glycerol to enhance wetting and hydraulic activation of stomatal pores. Within a few hours, nPAA-MnO_2_ accumulated in the sub-stomatal cavity and mesophyll apoplast, gradually releasing bioavailable Mn ions in the acidic apoplast environment. Moreover, labeling experiments with tracer ions revealed nPAA-MnO_2_ hot-spots around the vascular bundles and a limited but significant basipetal translocation of intact nanoparticles out of the foliar application zone, a pivotal step towards converting immobile nutrients such as Mn into mobile ones. Importantly, and unlike ionic Mn solutions, nPAA-MnO_2_ could be applied at high doses without causing leaf scorching and cytotoxicity, paving the way for more sustainable and efficient foliar fertilization practices. These novel aspects of nanoparticle uptake, translocation, and assimilation underscores the potential of nanotechnology to address nutrient mobility challenges in agriculture, representing an important contribution to the green transition of modern crop production.

## 1. Introduction

Mn is an essential plant micronutrient serving key functions in several metabolic processes such as photosynthesis, lignin biosynthesis and scavenging of reactive oxygen species.^1^ Mn is taken up from the soil in its divalent form Mn^2+^. Despite being abundant in the soil, Mn availability in the soil solution is heavily regulated by pH and redox status. Especially in calcareous, porous and sandy soils, Mn^2+^ ions are rapidly oxidized into Mn(IV)oxides becoming unavailable to plants. In a wide range of world crops, prolonged Mn starvation negatively impacts photosynthesis, increases plant sensitivity to oxidative stress, decreases water use efficiency and root functionality, which ultimately leads to substantial yield reductions.^2^ Under these circumstances, Mn is commonly applied as foliar sprays to crops to prevent or correct Mn deficiency. Foliar-applied Mn is rapidly taken up and assimilated by plants, resulting in a fast restoration of Mn metabolic functionalities in deficient plants.^3^ However, low phloem mobility of Mn prevents any significant translocation of Mn from the exposed tissue to the newly developing organs. Hence, several applications within a growing season are necessary to meet the Mn crop demand, representing a costly and wasteful practice for farmers. Furthermore, foliar application of Mn salts such as MnSO_4_ and MnCl_2_ can induce leaf scorching when applied in relatively high doses.

The introduction of nanotechnology in plant nutrition holds great promise in delivering alternative strategies for foliar fertilization, enabling more sustainable and cost-effective approaches. Comparably bigger than their ionic counterparts, yet small enough to penetrate all the relevant plant barriers e.g. cuticle, stomata, cell wall (CW) and vasculature, NPs can potentially deliver plant mineral nutrients right where needed, bypassing the limitations of ion mobility and enhancing nutrient use efficiency.^4^ Nanomaterials with different compositions, aspect-ratio, charge, and even size exceeding the size exclusion limit of the phloem barriers, have been reported to be taken up and translocated within the plant.^5–8^ Even though the mechanisms underlying NP remobilization remain to be clarified, the possibility of harnessing NPs’ ability to translocate represents a compelling argument for enhancing Mn foliar fertilization by adopting a nanotechnological approach. In addition, NPs may enable a slow and sustained release of nutrients inside the plant, relieving potential osmotic stress caused by overapplication of nutrient-containing salts onto the leaves, and thereby prevent scorching and tissue necrosis.

Reports on the effects of foliar application of Mn NPs are scarce,^9–13^ and the potential prospective benefits of Mn foliar nanofertilization are currently largely unexplored. Huang *et al.*^11^ reported a significant enhancement in wheat grains’ nutritional quality and yield following foliar treatment with MnFe_2_O_4_ NPs in comparison with ionic MnSO_4_ and FeSO_4_ combined. Similarly, Dimpka *et al.* ^9^ reported an increase in wheat shoot and grain Mn content, and greater grain yield after foliar exposure to nano-Mn, outperforming both nano-Mn and conventional Mn salts in soil applications. Other documented effects include greater photosynthetic efficiency, early-induced flowering and fruit production boost in tomato plants after foliar application of MnFe_2_O_4_.^13^ Finally, Elmer and White^10^ recorded enhanced resistance towards Fusarium wilt fungus in tomato plants sprayed with MnO NPs and grown in the pathogen-infested medium. Despite showing promising results, the aforementioned studies largely focus on agronomic performance, failing to provide an in-depth analysis of the mechanistic aspects regulating the interactions between Mn NPs and plants, and real NP uptake and utilization still awaits experimental documentation.

The objective of the current work is to study the effect of polymer-coated MnO_2_ NPs on plant growth and photosynthetic functionality of Mn-deficient barley plants, as well as provide mechanistic insights regarding uptake, distribution, translocation and metabolic assimilation of the foliar-applied NPs. We optimized the properties of the formulation to enable effective NP penetration through the leaf surface, securing the delivery of physiologically relevant amounts of bioavailable Mn, and we monitored the restoration of the Mn-dependent photosynthetic processes. Importantly, we employed a set of complementary techniques such as CLSM, nano-CT and LA-ICP-MS to reveal NP uptake pathways and allow the identification of NPs at the cellular level. In parallel, we used different analytical methods to validate NP presence and identity in plant tissue, and we quantified the uptake and distribution of NPs in young barley plants.

## 2. Results

### 2.1 Plant growth and Mn nutritional status

Control plants were Mn-sufficient throughout the 31 days of monitoring, with quantum yield of photosystem II (PSII) values (Fv/Fm) well above 0.75, indicating no restriction in electron flow between the photosystems (Figure **1A**). The nutritional status of Mn-deficient plants started declining to mild Mn-deficiency after 14 days and decreased further to moderate Mn-deficiency at 21 days with an Fv/Fm value of 0.55. Simultaneously, the typical visual symptoms of Mn deficiency started to manifest on the younger leaves, which appeared slack and showed interveinal chlorosis (Figures **1B, C**). Repetitive additions of Mn to the hydroponic solution after 21 days maintained Mn deficiency at a reversible stage, which allowed us to study how different foliar treatments were able to restore Mn functionality. Inductively coupled plasma optical emission spectroscopy (ICP-OES) data from plants at 21 days after transplantation (DAT) confirmed that Mn concentration in the youngest fully expanded leaf (YFEL) tissue dropped below the 15 μg g^-1^ dry matter (DM) (Figure **S1**), equivalent to the critical threshold for Mn deficiency.^2^

**Figure 1:**
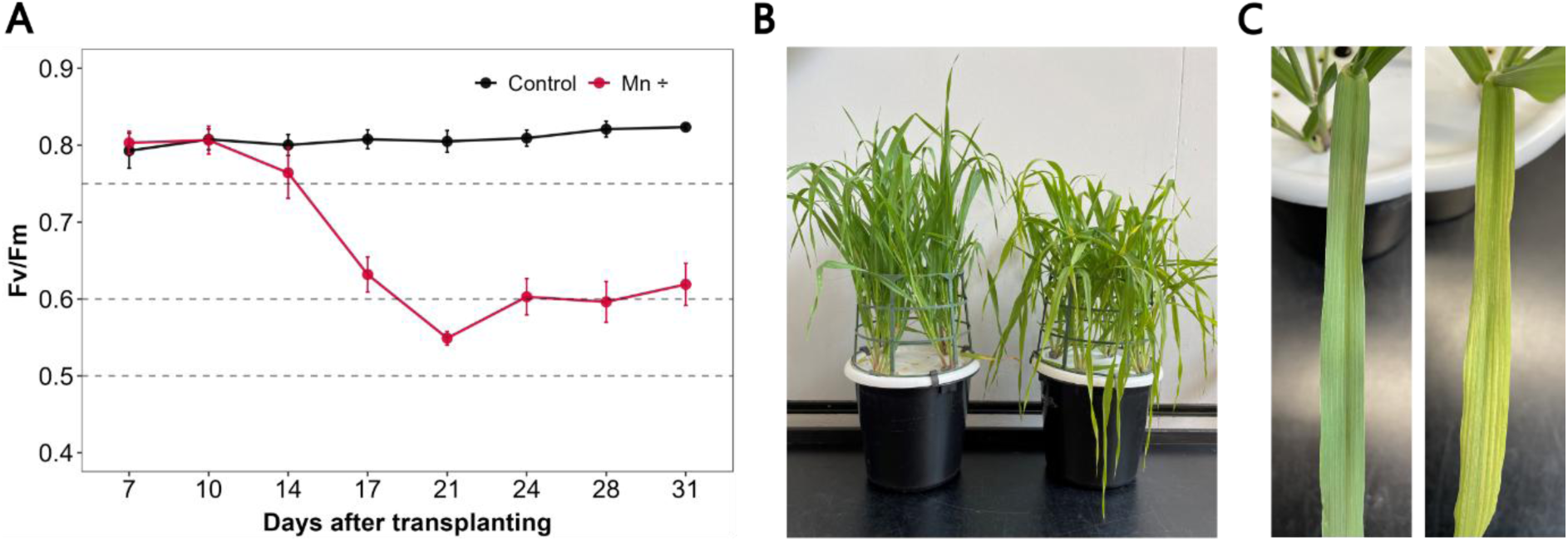
(A) The progression of Fv/Fm in YFEL indicating the plant Mn nutritional status in Mn-sufficient (*Control*) and Mn-deficient (*Mn ÷*) plants over time. Intercepts at 0.75, 0.6 and 0.5 represent the thresholds for mild, moderate and severe Mn-deficiency, respectively. Error bars show SD (n=4 with 3 technical replicates). (B, C) Photos of 21 DAT control plants (*left*) and plants with moderate Mn deficiency (*right*) displaying the characteristic interveinal chlorosis.

### 2.2. Nanoparticle synthesis and characterization

Transmission electron microscopy (TEM) and atomic force microscopy (AFM), employed to determine the morphology of pristine nPAA-MnO_2_, showed that NPs possessed a globular shape with a core of approximately 8 nm (Figures **2B, C** and **S2**). Dynamic light scattering (DLS) measurements indicated an average hydrodynamic diameter of 25.9 ± 5.3 and zeta potential of -46.4 ± 3.7mV (Table **S1**). The NP composition was analyzed by thermogravimetric analysis (TGA), showing a high content of PAA, which constitutes around 50% of nPAA-MnO_2_ mass (Figure **S3**). X-ray powder diffraction (XRD) pattern of a powdered nPAA-MnO_2_ revealed that the sample consists of the layered δ-MnO_2_ (birnessite) phase (Figure **S4**).^14^. The detection of a non-negligible amount of Na in the sample by inductively coupled plasma mass spectrometry (ICP-MS) (molar ratio Na:Mn of 1:2.6) is consistent with this finding, as Na^+^ is often found intercalated between the MnO_6_ octahedra, occupying the vacancies in the birnessite lattice.^15^ A distinct Na peak was also visible in the spectrum collected by transmission electron microscopy with energy dispersive X-ray spectroscopy (TEM-EDX) (Figure **S5**). Raman spectrum recorded on a nPAA-MnO_2_ liquid suspension showed strong peaks at 637 cm^-1^ and 580 cm^-1^, which are typically attributed to the Mn–O stretching vibration of the MnO_6_ octahedra and the basal plane of the MnO_6_ sheet of birnessite, respectively (Figure **S6**).^16,17^ Furthermore, the Raman band at 637 cm^-1^ could be assigned to the ν_1_-stretching mode of Mn^4+^O_6_, while the Raman band at 580 cm^-1^ to the ν_1_-stretching mode of Mn^3+^O_6_, indicating the Mn^4+^ and Mn^3+^ are the dominant oxidation states of Mn in nPAA-MnO_2_.^18^

**Figure 2:**
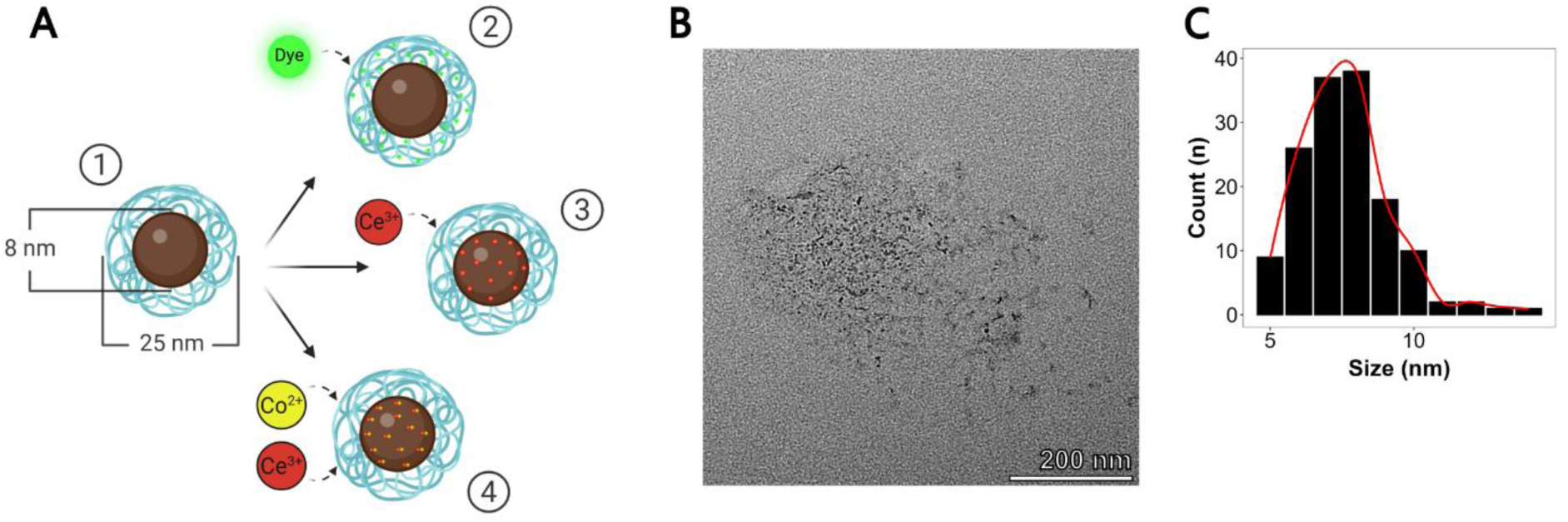
Size and morphology of of nPAA-MnO_2_. (A) schematic representation of the nPAA-MnO_2_ synthesized for the study, namely: (1) pristine nPAA-MnO_2_, (2) DiI-labelled nPAA-MnO_2_ (DiI-nPAA-MnO_2_), (3) cerium (Ce) spiked nPAA-MnO_2_ (Ce-nPAA-MnO_2_), (4) double-spiked Ce and cobalt (Co) nPAA-MnO_2_ (Ce/Co-nPAA-MnO_2_). (B) TEM image of nPAA-MnO_2_. (C) Size distribution of the nPAA-MnO_2_ core from figure B determined using ImageJ. Size refers to the Feret’s diameter (n=144).

Ce-nPAA-MnO_2_ and Ce/Co-nPAA-MnO_2_ morphology, hydrodynamic diameter and zeta potential are consistent with the pristine nPAA-MnO_2_, suggesting no changes to their main physical properties (Figure **S7** and table **S1**). Elemental analysis by ICP-MS of Ce-nPAA-MnO_2_ and Ce/Co-nPAA-MnO_2_ was used to evaluate quantify the incorporation of tracer elements into the NP structure. Ce-nPAA-MnO_2_ possessed a Ce:Mn molar ratio of 1:99, whereas in the double-spiked Ce/Co-nPAA-MnO_2_ the molar ratios detected for Ce:Co:Mn were 1:1:99, close to the 1:100 intended ratio between Mn and the tracer elements. The efficient incorporation of cationic tracers into the NP structure is likely facilitated by the strong cation sorption capacity of MnO_6_ octahedra of birnessite due to the large number of cation vacant sites.^14,19,20^

The fluorescence spectroscopy analysis confirmed the successful encapsulation of the DiI dye, as a red shift in the DiI-nPAA-MnO_2_ (588 nm) emission spectrum was observed as opposed to DiI alone (580nm) (Figure **S8B**). Furthermore, no changes in zeta potential and hydrodynamic diameter before and after labelling were measured by DLS, indicating that the original properties were preserved (Figure **S8C**).

### 2.3. Formulation of nanoparticles

*In vitro* dissolution tests showed a pH-dependent dissolution profile for the nPAA-MnO_2_ (Figure **S9**). At pH 5.5, the dissolution was the highest, with 99% of Mn released from the NPs within 3 days. The inverse relationship between pH and Mn release is showcased at increasing pH values: at pH 6.5 and 7.5, 71% and 20% of Mn was liberated after 3 days, respectively.

The surface tension of the different solutions (*i.e.* water, MnSO_4_, NPs and NPs with glycerol) is shown in figure **S10**. MnSO_4_ and NPs solutions had surface tension values above 70 mN m^-1^, comparable to ultrapure H_2_O, whereas the NPs solution with 3% glycerol had lower surface tension of ∼66 mN m^-1^. The surface tension of formulations at varying concentrations of Silwet Gold surfactant revealed a clear trend of decreasing surface tension with increasing surfactant concentration. A similar response was obtained for all the different formulations and the 0.1‰ concentration of Silwet Gold yielded surface tension values at or slightly lower than 20 mN m^-1^.

### 2.4. Foliar application and restoration of Mn functionality in deficient barley plants

To assess the physiological plant response to different Mn sources and concentrations, we tested MnCl_2_, MnSO_4_ and nPAA-MnO_2_ solutions on Mn-deficient barley plants. No effects were observed when the formulation alone (*Control*: 0.1‰ Silwet Gold, 3% glycerol) was applied (Figure **3A**). On the contrary, scorching symptoms and necrotic spots were visible three days after the application of MnCl_2_ and MnSO_4_ solutions in agronomic relevant concentrations. Also, the scorching intensity was concentration-dependent and appeared more severe for MnCl_2_. Furthermore, necrotic bands on the treated leaves were visible beyond the droplet deposition area towards the leaf tip. Conversely, no scorching was reported for the nPAA-MnO_2_ treatment when applied in similar concentrations.

**Figure 3:**
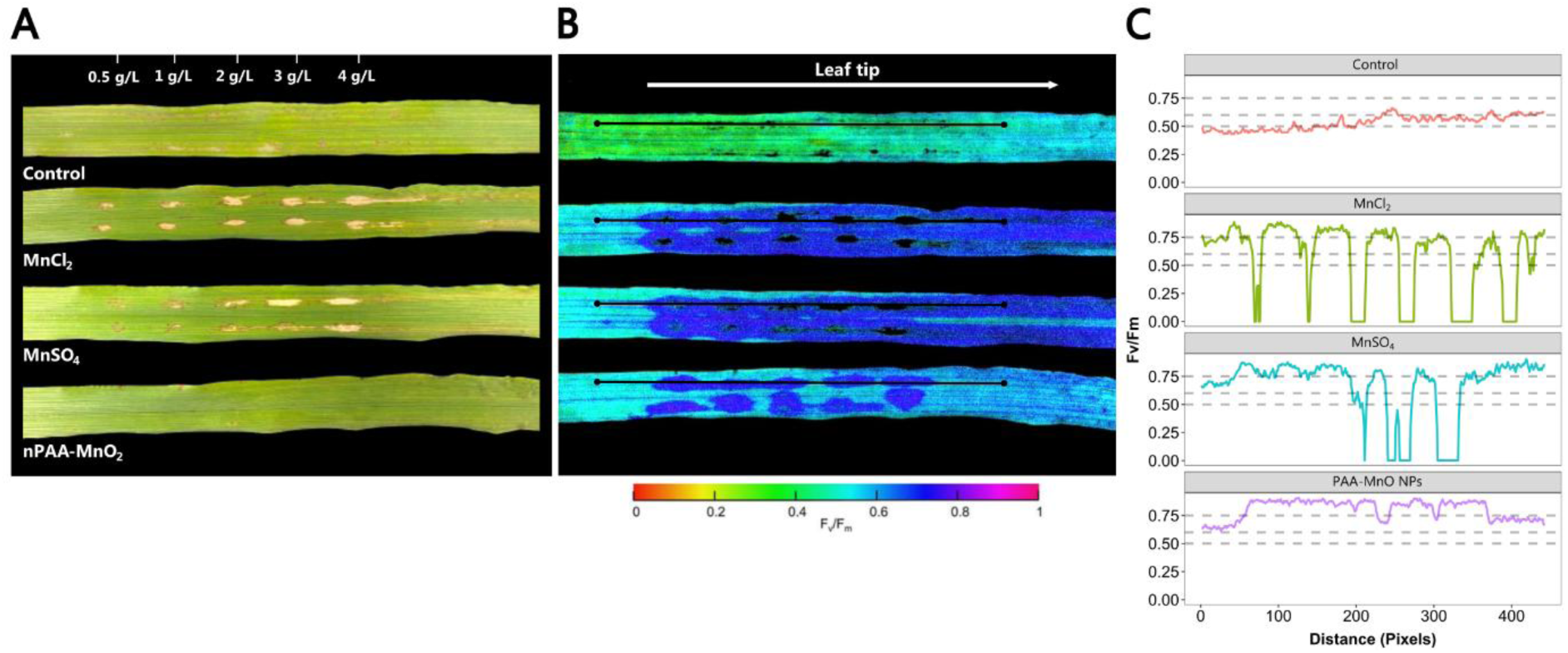
(A) representative photo of Mn-deficient barley leaves three days after exposure to droplets of different Mn sources and concentrations. From top to bottom: control (0.1‰ Silwet gold, 3% glycerol), MnCl_2_, MnSO_4_ and MnO NPs. The numbers above indicate the concentration (g/L) of Mn contained in the different solutions except for the control, which did not contain any Mn. (B) Corresponding color-coded pulse amplitude modulation (PAM) fluorometry image showing the spatial Mn-status distribution within the treated leaves. Colored scale bar shows Fv/Fm values from 0 to 1. The blue color of the scale indicates higher Fv/Fm values, typical of Mn-sufficient plant tissues. Transects (black lines) were drawn through the leaf treated area. (C) The curves illustrate the Fv/Fm values measured by transects. Horizontal dashed lines indicate mild (0.75–0.6), moderate (0.6–0.5) or severe (<0.5) Mn deficiency.

Subsequently, to verify the effects on Mn dependent photosynthetic functionality, the treated leaves were analyzed using imaging PAM (Figure **3B**). The leaves that received droplets of the MnCl_2_ and MnSO_4_ solutions showed a broad Fv/Fm restoration area extending towards the leaf tip, whereas in the leaf dosed with nPAA-MnO_2_ the Fv/Fm restoration was confined to the areas where droplets where deposited. Finally, transects showed that all Mn sources promoted Fv/Fm restoration to control levels (Fv/Fm > 0.75; Figure **3C**). However, Fv/Fm values of 0 were recorded in the necrotic areas where MnCl_2_ and MnSO_4_ droplets were applied.

### 2.5. Uptake efficiency of nanoparticles

The effects of glycerol addition on the uptake efficiency of NP formulations (*NPs (S+G)* vs *NPs (S)*) was studied in Mn-deficient plants using Ce-nPAA-MnO_2_ and compared against an ionic treatment containing MnSO_4_ and CeSO_4_ (*Ionic (S+G)*) (Figure **4**). We quantified the uptake of the different foliar formulations by elemental analysis in barley YFELs 3 days after foliar exposure. As Mn is an essential nutrient to plants, a relatively high Mn background is found in tissues. Thus, Ce which virtually has a zero background in plants, was embedded into the nPAA-MnO_2_ structure to trace their uptake.

**Figure 4:**
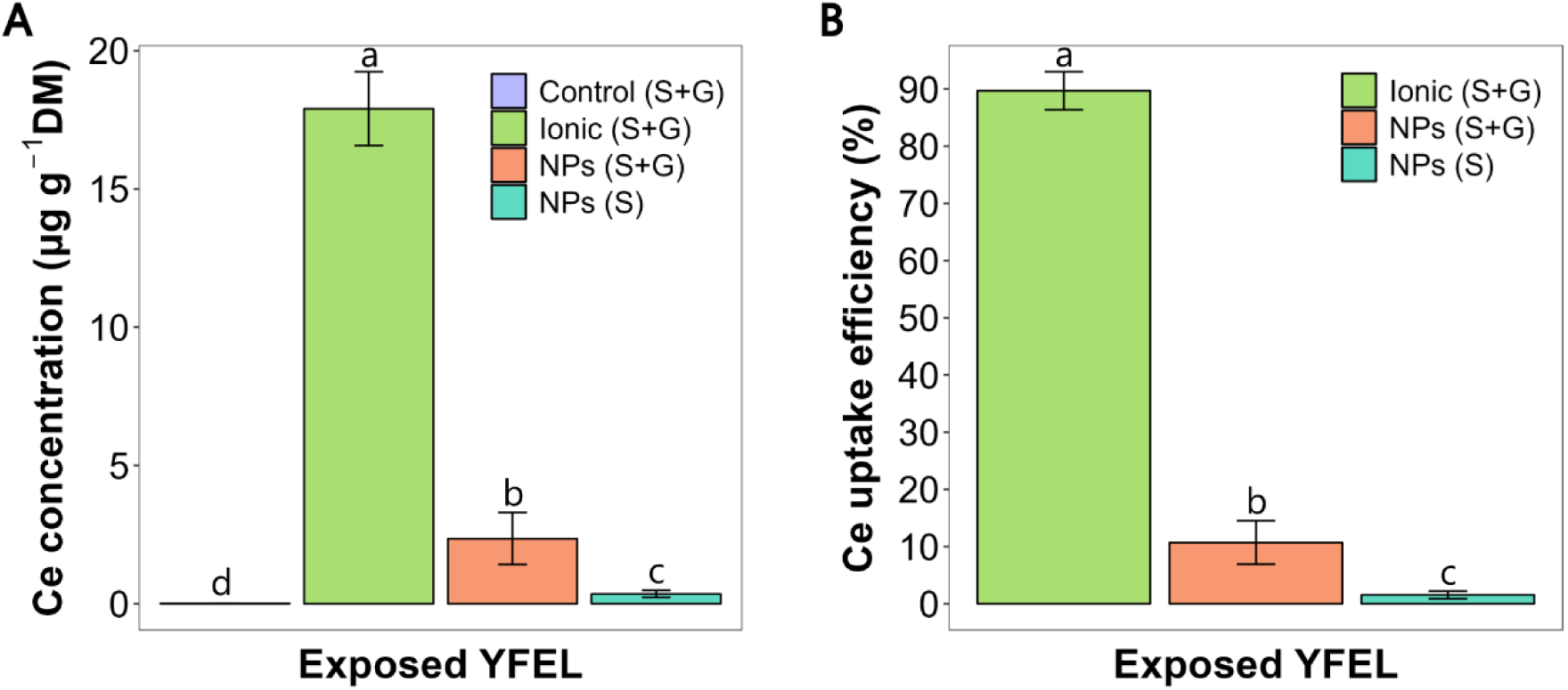
Uptake efficiency of foliar-applied ions versus NPs, formulated with or without 3% glycerol. YFELs were sampled 3 days after NP exposure. (A) Ce concentration in the exposed YFELs. (B) Uptake efficiency of Ce in the different treatments measured as a percentage of the total amount applied. See Material and Methods for details. Results are presented as means ± SD (n = 6), and letters represent significant differences (p < 0.05) analyzed by a one-way ANOVA and Tukey’s multiple comparison test.

Following application, the concentration of Ce tracer in the exposed YFELs was significantly higher than the control for all the treatments (Figure **4A**). After converting the Ce concentration in exposed YFELs into uptake efficiency, we observed an uptake of 90%, 11%, 2% (P < 0.05) in the ionic (S+G), NPs (S+G) and NPs (S) treatments, respectively (Figure **4B**). Similar patterns were reported for Mn (Figures **S11A**, **B**), for which we observed uptake rates of 79%, 9%, 2% (P < 0.05), in the ionic (S+G), NPs (S+G) and NPs (S) treatments, respectively.

Glycerol presence in the formulation extended the drying time of droplets with up to 3 hours after deposition, in contrast to the NP solution containing Silwet Gold only, which dried up within 1 hour (Figure **S12**).

### 2.6. Mechanistic aspects of foliar nanoparticle uptake

We investigated foliar uptake and mesophyll distribution of nPAA-MnO_2_ by CLSM, nano-CT and LA-ICP-MS. In all cases, NPs were formulated with 0.1‰ Silwet Gold and 3% glycerol and dosed to the adaxial leaf surface. The samples used in these analyses were either directly imaged or embedded into a sample mounting medium between 2 and 5 hours after treatment. After 2 hours of nPAA-MnO_2_ exposure, confocal analysis showed that fluorescence localized to the sub-stomatal cavity along parallel sets of stomata (Figure **5**; Video **S1**). Inside the leaf, NPs were distributed in the apoplast surrounding mesophyll cells, with no sign of cell internalization. In contrast to the nPAA-MnO_2_ treatment, a different pattern was observed for the DiI-only formulation, which did not penetrate through the stomata over a period of 5 hours of incubation (Figure **S13**; Video **S2**). In this case, the DiI dye was found evenly spread on the leaf surface and on top of the stomatal aperture, but never in the sub-stomatal cavity.

**Figure 5:**
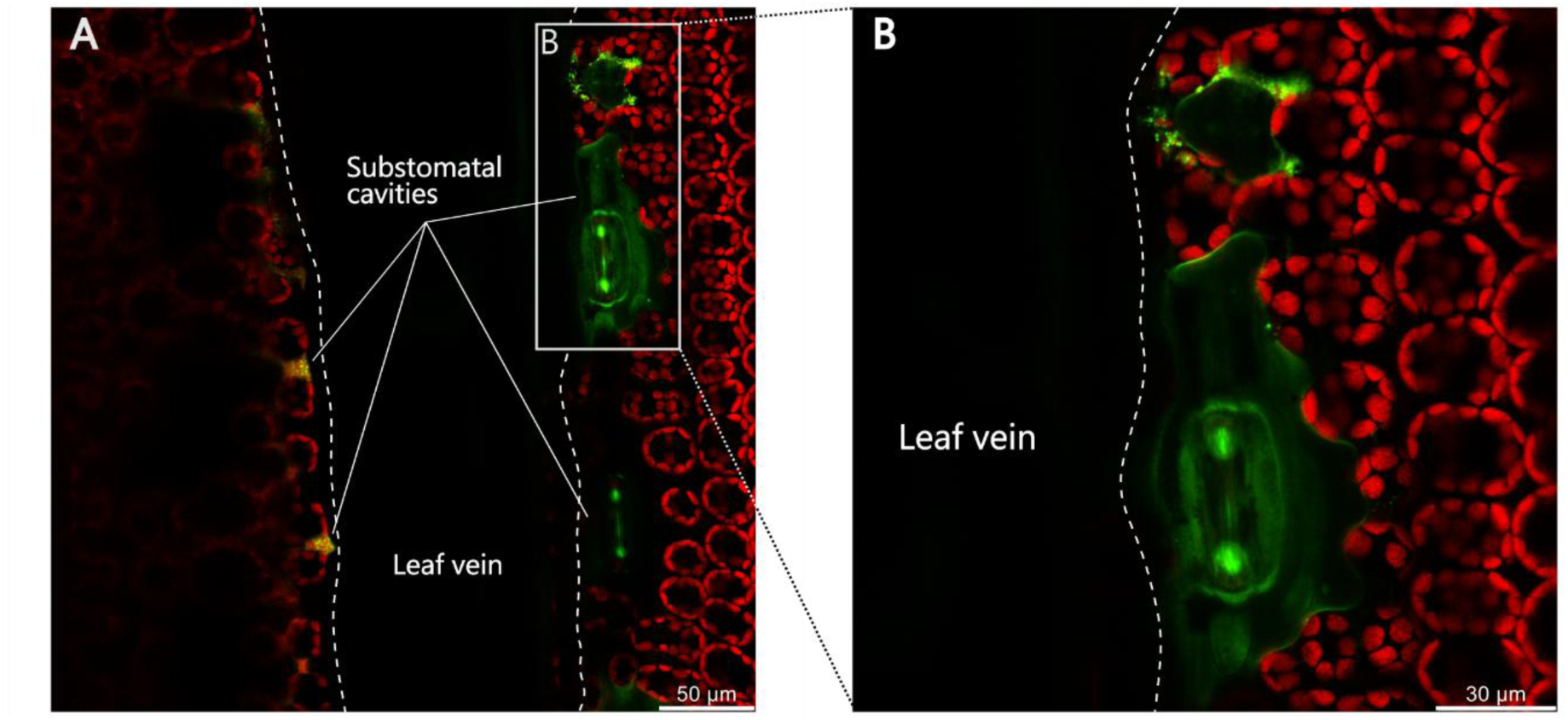
Confocal images of fluorescent DiI-nPAA-MnO_2_ in Mn-deficient barley leaves. (A) Top view of the NP treated leaf and (B) close-up of the highlighted area. In red: chlorophyll autofluorescence from the chloroplasts; in green: DiI signal.

To validate the findings from confocal imaging, we tested the capability of nano-CT for studying NP uptake pathways in leaves. Discrete nPAA-MnO_2_ were not visible when infiltrated in the leaf (Videos **S3**, **S4**). However, NP clusters were detected when NPs were sequentially infiltrated after 20 mM CaCl_2_, to promote NP aggregation inside the leaf (Videos **S5, S6**). To verify the presence of NP in the mesophyll apoplast, 20 mM CaCl_2_ was infiltrated prior to the topically application of NPs on the adaxial page. Following leaf application, small NP clusters became apparent in the mesophyll area below the stomata (Figure **6**; Video **S7**). CaCl_2_ infiltrated leaves were analyzed as controls and did not show any artefacts resembling NP clusters inside the mesophyll (Video **S8**).

**Figure 6:**
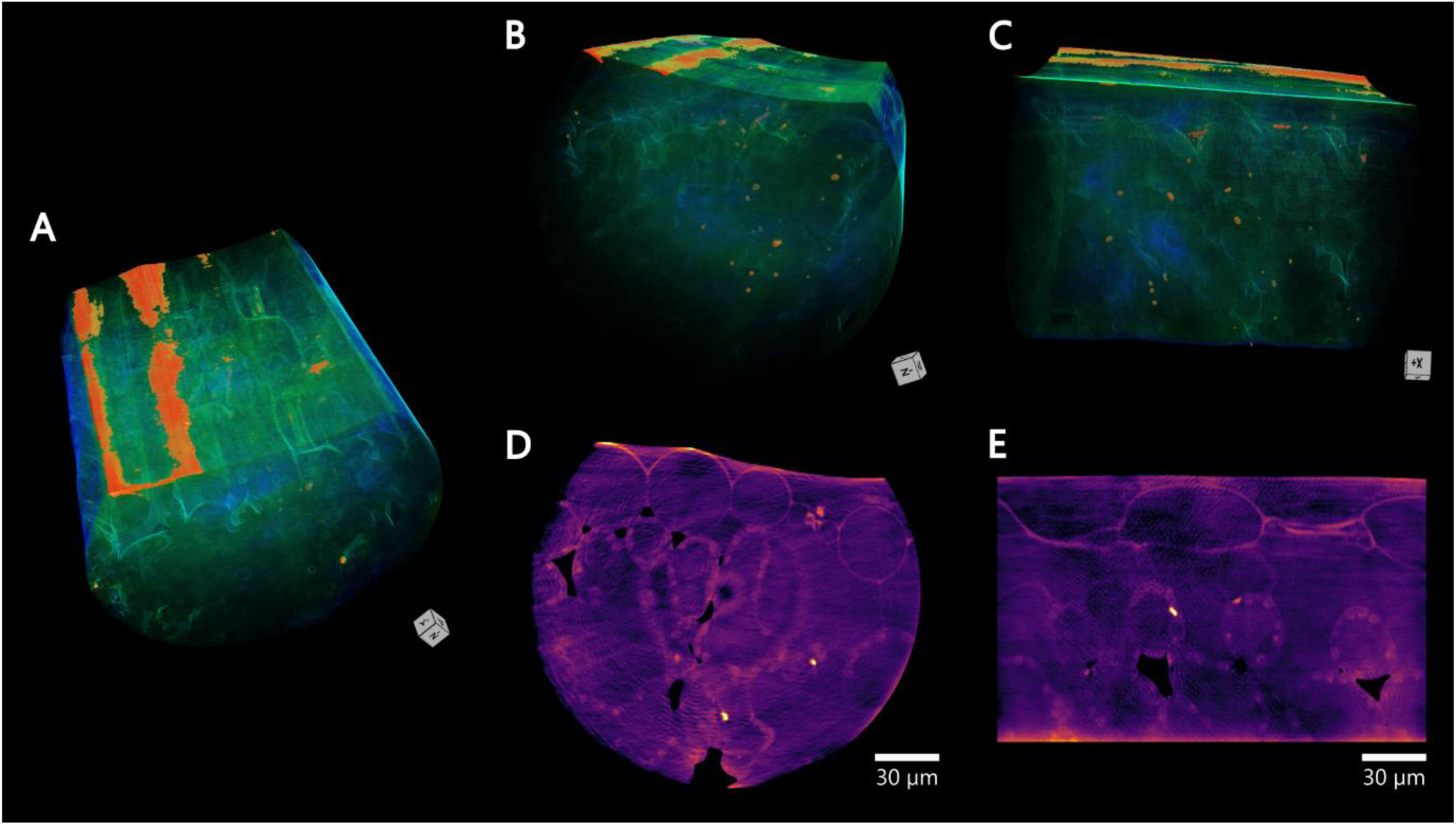
High resolution nano-CT images of a fresh barley leaf treated with nPAA-MnO_2_ following CaCl_2_ infiltration (pixel size 100 nm). Top (A), front (B) and lateral (C) views of the 3D leaf reconstruction showing NP clusters in red, leaf tissues in green and water in blue. NPs clusters are visible on the leaf surface (A) and in the mesophyll regions close to stomata (B and C). 2D virtual slices corresponding to the front (D) and lateral (E) views, showing NPs clusters as bright spots. The 3D leaf reconstruction can be viewed in video **S7**.

Finally, the nPAA-MnO_2_ distribution patterns in tissue were determined by LA-ICP-MS and correlated with those observed *via* confocal imaging and nano-CT (Figure **7**). LA-ICP-MS on barley leaf cross section highlighted different uptake pathways between Ce-nPAA-MnO_2_ and free Ce and Mn ions. Following the application of Ce-nPAA-MnO_2_, Mn and Ce signals strongly co-localized in proximity of sub-stomatal regions, across a group of fiber cells. A different uptake pattern was observed for ionic Mn and Ce, which appeared to permeate through the leaf surface as their signals were found around fiber cells, epidermal cells and close to stomata.

**Figure 7:**
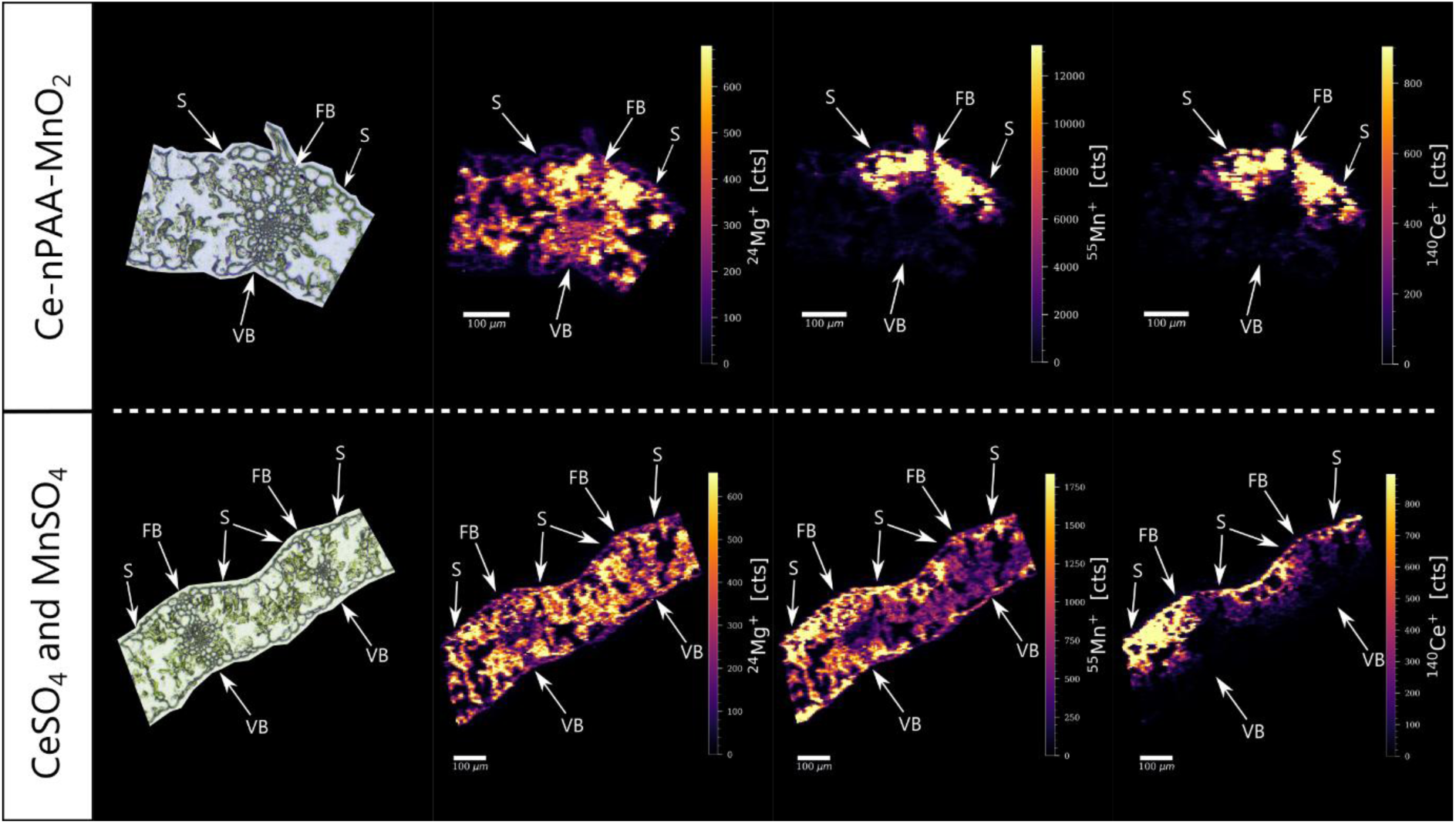
Elemental maps of 14 µm-thick barley leaf cross sections treated with Ce-MnO NPs and an ionic Mn and Ce solution. Mg is ubiquitously present in leaf tissue as a co-factor in chlorophyll and it was used to highlight the mesophyll. Laser spotsize 5 µm. The annotations indicate: S, stoma; FB, fiber cells; VB, vascular bundle.

The multi-elemental imaging showed that foliar-applied NPs and free ions distributed differently within the leaf (Figure **8**). Leaves treated with Ce-nPAA-MnO_2_ showed a strong Mn signal intensity in the mesophyll tissue, with clear hotspots corresponding to the vascular bundles displaying the highest ion intensities. An analogous distribution was observed for the NP tracer Ce, which was found as hot spots co-localizing with the leaf vasculatures (Figures **8A, B**). In contrast to NPs, the highest ion intensities for both ionic Mn and Ce were found in the mesophyll tissue, although they were also present in most leaf vascular bundles (Figures **8C, D**). Notably, the ionic Ce signal was mostly present in the upper mesophyll region, whereas the Mn was found spreading across the whole mesophyll. Mn-deficient control (untreated) leaf sections showed that native Mn was primarily associated with the mesophyll cells, with no or very little presence in the vascular bundle, bundle sheath, fiber cells and epidermis (Figure **S14**). Ce instead was not present in untreated leaves.

**Figure 8:**
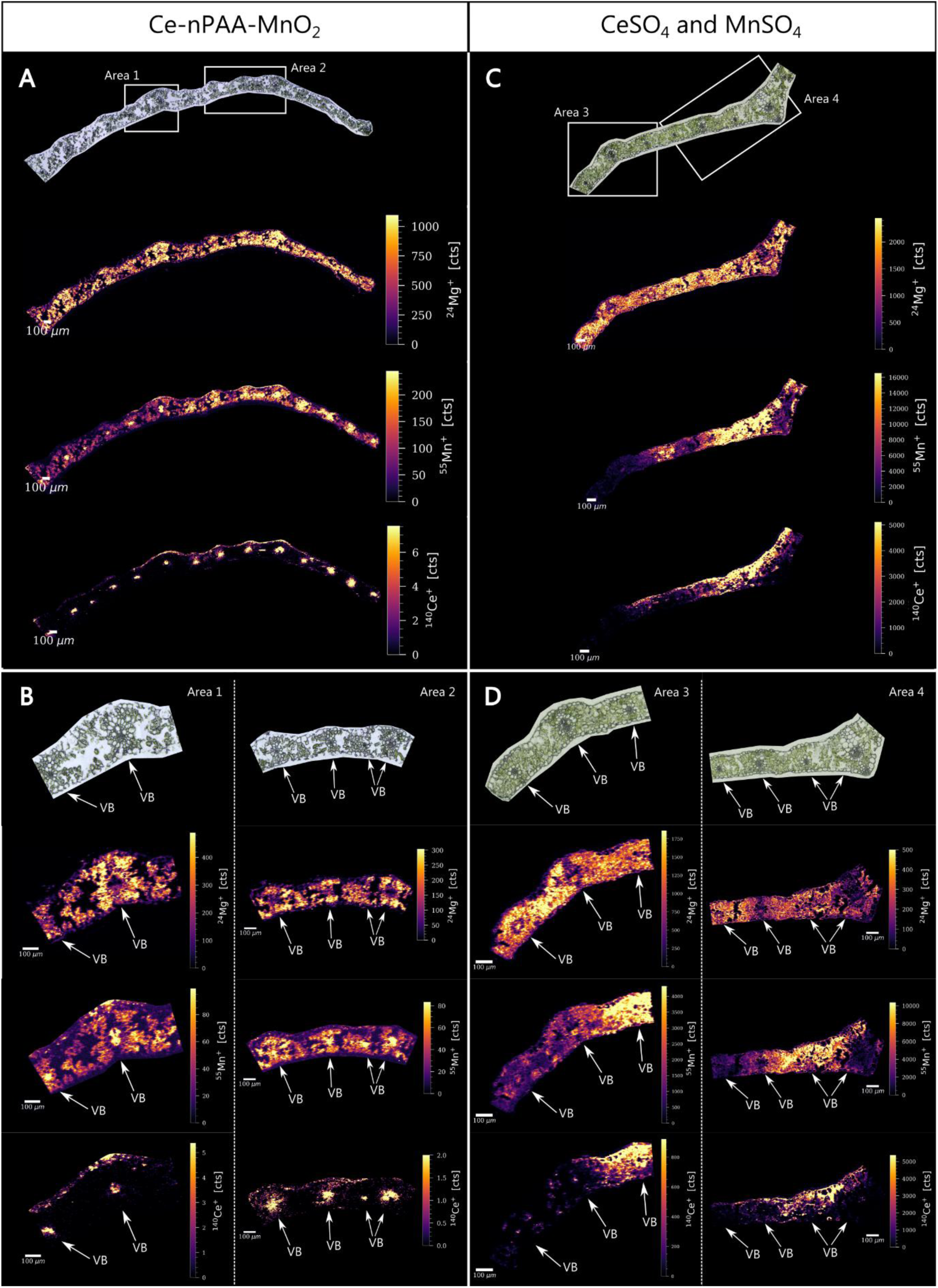
Elemental maps of 14 µm-thick barley leaf cross sections treated with Ce-MnO NPs and an ionic Mn and Ce solution. Mg is ubiquitously present in plant tissues and it was used to highlight the plant CWs. (A) Overview of the Ce-PAA-MnO NPs treatment (spotsize 5 µm). (B) Vasculature close-ups of the areas 1 and 2 (spotsize 4 µm). (C) Overview of the leaf treated with a MnSO_4_ and CeSO_4_ solution (spotsize 5 µm). (D) Vasculature close-ups of the areas 3 and 4 (spotsize 4 µm). Annotation indicates: VB, vascular bundle.

### 2.7. Assimilation and distribution of Ce/Co-nPAA-MnO_2_ nanoparticles in plants

Modified Ce/Co-nPAA-MnO_2_ were used to study *in planta* dissolution and translocation of NPs by elemental analysis of Ce and Co in barley leaves, shoots and roots. In addition, the NP treatment was compared against an ionic formulation containing MnSO_4_, CeSO_4_ and CoCl_2_. Owing to different mobility in plants, phloem immobile Ce and highly mobile Co ions were considered as proxies for nPAA-MnO_2_ remobilization and dissolution, respectively.

Prior to Ce/Co-nPAA-MnO_2_ testing on plants, the release kinetics of the NP core elements (Mn, Ce and Co) were determined in a pH 5.5 citrate buffer resembling the pH of the apoplast and cell wall environment. The dissolution assay showed that Mn and Co were released at a similar rate, as their molar ratio (1:100) was nearly preserved over time (Figure **S15A**), while trivalent Ce ions was released at a lower rate compared to divalent Mn and Co (Figures **S15B**, **C**). The slower Ce release in the first 24 hours of the dissolution assay resulted in higher Mn:Ce and lower Ce:Co molar ratios (Figures **S15B**, **C**).

Four days after the application of foliar solutions, symplastic Ce was mostly found in the exposed leaf and the concentration was highest for the ionic treatment (Figure **9B**). Examining the Ce distribution pattern beyond the exposed area, the only statistically significant differences were found in the “Adjacent” fraction, where the Ce concentration for both the ionic and NP treatments were significantly higher that the untreated control. After computing the percentage of the Ce content in the “Adjacent” fraction relative to the Ce content in the “Exposed” area, it was calculated that 0.02% and 0.26% Ce (P < 0.05) was translocated in basipetal direction from the application area in the ionic and the NP treatment, respectively. Similarly to Ce, Co was found accumulating in the exposed area and again the uptake was higher for the ionic treatment compared to NPs (Figure **9C**). Relatively small, but significant enrichments of Co, were detected in the “Adjacent” and “Shoot” fractions for both the ionic and NPs treatments. However, the presence of Co in the roots was observed only for the ionic treatment, underlining the high phloem mobility of Co ions. Looking at the Mn distribution, this was largely found in the “Exposed” fraction, with the greatest Mn enrichment reported for the ionic treatment (Figure **S16**). Beyond the exposed area, significant increases in Mn concentration were detected in the “Adjacent” fraction for both the ionic and the NP treatment. After calculating the amount of Mn in the “Adjacent” as a percentage of the “Exposed” content, it was found that 0.14% and 1.93% (P < 0.05) of Mn translocated in the basipetal direction from the exposed to the adjacent area in the ionic and the NP treatment, respectively. No differences were found in the shoot and roots (Figure **S11**).

**Figure 9:**
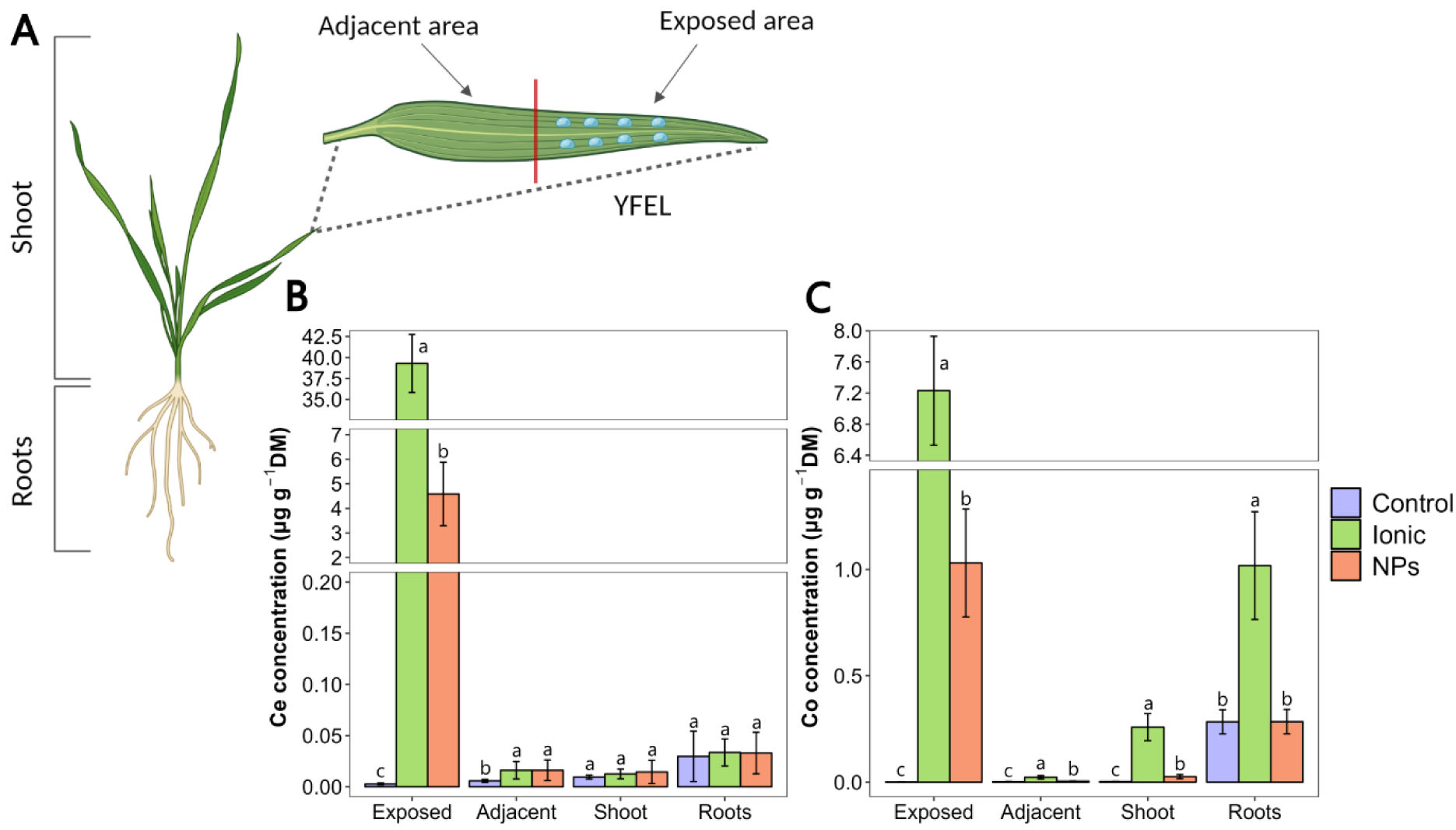
Symplastic Ce and Co distribution in different plant fractions 4 days after foliar exposure to Ce/Co-nPAA-MnO_2_ and an analogous ionic solution. (A) Schematic representation of the foliar treatment setup including the sampled fractions: exposed area, adjacent area, shoot and roots. (B) and (C) show the Ce and Co distributions in the different plant fractions, respectively. The legend shows the three different treatment consisting of different formulations, namely: Silwet Gold and glycerol (*Control*), MnSO_4_, CeSO_4_ and CoCl_2_ (*Ionic*), and Ce/Co-nPAA-MnO_2_ (*NPs*). Results are presented as means ± standard deviation of the mean (n = 6), and letters represent significant differences (p < 0.05) analyzed by a one-way ANOVA and Tukey’s multiple comparison test carried out within groups.

## 3. Discussion

### 3.1. Restoration of Mn functionality

The imaging-PAM assay (Figure **3**) showed that application of nPAA-MnO_2_ droplets promoted a full restoration of PSII functionality in Mn-deficient leaves within 3 days after foliar treatment. Similarly, ionic MnCl_2_ and MnSO_4_ solutions induced full restoration as indicated by the quantum yield of photosystem II. However, when applied in the same Mn concentrations, Mn salts caused severe leaf scorching, which appeared to differ according to the counter ion, possibly owing to differences in point of deliquescence (POD) between chloride and sulfate salts.^21^ The lower POD of chlorides would maintain MnCl_2_ in solution for a longer period compared to MnSO_4_, promoting its leaf absorption and thereby resulting in greater osmotic and phytotoxic effects. Conversely, nPAA-MnO_2_ did not cause any phytotoxicity even at high concentrations.

Once inside the leaf, nPAA-MnO_2_ released Mn^2+^ gradually in response to the slight acidity of the apoplast as demonstrated under *in vitro* conditions (Figure **S9**). Slow Mn^2+^ release combined with the lower foliar uptake of nPAA-MnO_2_ in comparison to the ionic forms are the probable causes for the absence of phytotoxic effects on leaves following nPAA-MnO_2_ application. Despite the leaf scorching, droplets of soluble MnCl_2_ and MnSO_4_ were more efficient at restoring Mn deficiency. This was mainly caused by the observation that Mn^2+^ from these salts were translocated in acropetal direction out of the labeling zone with the transpiration stream, spreading towards the leaf tip and margin, thus promoting recovery of Mn-functionality in a broader leaf area compared with the nPAA-MnO_2_.

Using the same Mn concentrations, nPAA-MnO_2_ showed little tissue spreading and performed less effectively when applied as larger droplets than ionic forms over the experimental period considered (3 days). Nonetheless, our results clearly show that NPs can provide adequate amounts of bioavailable Mn to deficient plants, whilst basically eliminating the risks associated with excessive nutrient application such as phytotoxicity. To date, no studies had clearly shown the benefits of the foliar application of Mn containing NPs compared to their ionic counterparts. In most cases, studies were conducted on plants where no Mn-nutrient limitation was reported, the amounts of applied Mn were not specified, and more importantly, no real leaf uptake of intact NPs was documented.^9–12^. Notably, our findings demonstrate the effectiveness of Mn containing NPs in restoring essential functions in plants and their potential to support fundamental metabolic processes by supplying a pH responsive and sustained release of mineral ions.

### 3.2. Formulation and uptake efficiency

The use of adjuvants is key to ensure proper contact of foliar sprays with the leaf surface and promote the penetration of agrochemicals. Yet, the role and impact of formulation properties on NP leaf uptake are generally overlooked. The results presented in this study show that a combination of surfactant and humectant was crucial to promote NP uptake in barley leaves (Figure **4**). nPAA-MnO_2_ formulated with 0.1‰ Silwet Gold and 3% glycerol were taken up 5 times better (Figure **4B**) compared to a formulation containing Silwet Gold only. While a low surface tension (∼20 mN m^-1^) is required to promote penetration of foliar formulations,^22^ the addition of a humectant such as glycerol kept the droplets moist for longer (Figure **S12**), likely maintaining the NPs in suspension and allowing them to diffuse through the water film continuum connecting the leaf surface to the interior. PAA constitutes ∼50% of the nPAA-MnO_2_ mass (Figure **S3**), and while being a hydrophilic superabsorbent polymer, it most likely acted synergistically with glycerol contributing to retain water molecules thereby preserving a thin water film for an extended period. To our knowledge, only Hu *et al.* ^23^ previously reported on the use of glycerol in a NP formulation. Similarly, they observed leaf NP uptake in maize only when 3% glycerol was added to a formulation containing Silwet-77 surfactant. In contrast, no humectant was required to promote leaf NP uptake in cotton leaves, highlighting key differences in response across different plant species. Remarkably, leaf uptake has also been reported for foliar-applied NPs prepared with no adjuvants.^5^ Studying the leaf interaction and uptake dynamics of 3, 10 and 50 nm citrate- and PVP-coated AuNPs in wheat plants, Avellan *et al.* (2019) found that the smaller NPs (*i.e.* 3 and 10 nm AuNPs) were strongly retained on the leaf surface, and that amphiphilic PVP coating promoted efficient leaf NP uptake *via* the cuticle. This shows that NP surface functionalities and size are additional key players in regulating NP leaf adhesion and uptake processes.

Overall, leaf NP interactions are governed by several factors including formulation parameters, NP physicochemical properties,^5^ plant anatomical and morphological features^23^ and environmental factors.^4^ Further research should investigate the role of isolated variables on the uptake efficiency of foliar-applied NPs across different crop species and investigate how *e.g.* humidity levels can affect cuticular pores and stomatal hydration and thus NP uptake.

### 3.3. Mechanisms of nanoparticle uptake and leaf tissue distribution

Complementary CLSM, nano-CT and LA-ICP-MS provided evidence that foliar-applied nPAA-MnO_2_ NPs rapidly entered the leaf *via* stomata (Figures **5**, **6**, **7**). Interestingly, LA-ICP-MS highlighted a different uptake pathway for ionic Ce and Mn, which appeared to enter the leaf *via* small hydrophilic cuticular pores, as their signals were uniformly found around epidermal cells and fiber cells, as well as through stomata (Figure **7**). Mineral ions have been previously reported to enter *via* the fiber cells and cuticle.^24^ It has been hypothesized that the hydration of the cuticular polar domains can promote swelling, resulting in the formation of larger pores which can facilitate the passage of solutes across the leaf surface.^24–26^ In this case, the presence of glycerol in the formulations used here, has likely contributed to prolonging the hydration of the leaf surface, activating the cuticular pathway and thereby promoting ion transport into the leaf.

Following uptake, Ce and Mn signals from the nPAA-MnO_2_ treatment strongly co-localized with the leaf vasculature, where clear hotspots were found (Figures **8A, B**). In contrast, the highest ion count rates for both ionic Mn and Ce were found throughout the mesophyll tissue, where they appeared to accumulate, although they were also present in most leaf vascular bundles (Figures **8C, D**). Such differences may be explained by distinct mechanisms that regulate ions and NPs transport within the leaf. Ion uptake into the mesophyll apoplast is controlled by diffusion and primarily driven by the concentration gradient.^3^ As for NPs, the forces controlling their movement within the leaf remain elusive. Yet, it is possible that phloem loading/unloading, xylem counter-flow and phloem/xylem exchange processes may promote NP transport towards the vasculature and the subsequent accumulation there, although efforts are warranted to address this major fundamental research gap.^27–29^

In this study, we attempted to bypass the limitations of individual techniques by employing a multimodal bioimaging approach to obtain a clear picture of leaf NP uptake and distribution pathways. For instance, the interpretation of the images obtained by CLSM may be complicated by the risk of fluorescent tags dissociating from the NPs or freely diffusing upon NP dissolution. Here we show that the foliar-applied DiI dye produces distinctively different leaf distribution patterns in comparison to when this is incorporated into the structure of nPAA-MnO_2_ and therefore it could be used as a control treatment in confocal microscopy (Figures **5**, **S13**; Videos **S1**, **S2**). DiI is a lipophilic dye used for staining cell membranes and other hydrophobic structures.^30^ Being a lipophilic molecule, DiI is expected to preferentially bind the abundant apolar moieties of the cuticle, which may prevent it from following the diffusive water film connecting the surface to the hydrophilic leaf interior.

The poor tissue penetration of light in confocal microscopy limits its use in the study of processes occurring a few hundred microns below the leaf surface. Thus, in the present study, we tested the capabilities of nano-CT to obtain high-resolution, three-dimensional images of label-free nPAA-MnO_2_ in fresh leaf samples. In agreement with CLSM analysis, nano-CT reconstructions highlighted the presence of NP clusters in the mesophyll region close to stomata following NP foliar application (Figure **6**; Video **S7**). However, nano-CT could not be used to visualize nPAA-MnO_2_ unless aggregation was induced by leaf infiltration with CaCl_2_. Interestingly, despite being forced to aggregate, small NP clusters were visible inside mesophyll cells in proximity to the cell wall (Figures **6D, E**). This evidence may suggest that nPAA-MnO_2_ aggregation occurred slowly and that NP clusters were formed following NP transport across the cell wall. NPs of similar size to nPAA-MnO_2_ have been reported to cross the CW.^23,31^ Previous studies have also demonstrated the ability of highly-charged PAA-coated CeO NPs to cross the CW and localize in the chloroplasts upon foliar exposure to NPs.^23,32^ However, given that NPs were forced to aggregate, it could be questioned as to whether nano-CT images are truly representing uptake and distribution dynamics of NPs in leaf tissue under native conditions. Undoubtedly, it is not possible to extrapolate information on how NPs may further distribute beyond the sub-stomatal cavity using nano-CT, as it is likely that the state of aggregation of NPs influences their transport within the leaf. Concomitant factors such as the size and colloidal stability of nPAA-MnO_2_ are critical features that may prevent their detection in tissue under normal conditions (*i.e.* without inducing aggregation). Highly charged nPAA-MnO_2_ have demonstrated remarkable stability in solution under a wide range of concentrations, possibly owing to the abundant carboxylic groups on the NP surface which provide electrostatic and steric repulsion preventing particle aggregation.^33^ The low propensity of NPs to aggregate could explain why NPs were not detected even upon leaf infiltration (Videos **S3**, **S4**). Additionally, small differences in electron density between the soft leaf tissue and the polymer-rich nPAA-MnO_2_ did not produce substantial contrast to distinguish single NPs from the plant tissue background. To date, only Avellan *et al.* ^34^ explored the capabilities of nano-CT imaging to study Au NPs interactions with plant roots. The authors detected large Au NPs agglomerates associated with the root cap in a dehydrated and dried sample. As also suggested by that study, NP aggregation appears to be a prerequisite for detection in nano-CT imaging. In fact, despite its high resolving power, the size of a single nPAA-MnO_2_ is below the nano-CT voxel size (*i.e.* 50 nm) and thus its visualization could not be expected unless some aggregation takes place.

To complement the morphological data obtained with CLSM and nano-CT, we integrated LA-ICP-MS to map the spatial distribution of Ce-nPAA-MnO_2_ at the whole leaf level (Figure **8**). Interestingly, different patterns were observed when Ce and Mn were applied in nanoparticulate or ionic forms. However, LA-ICP-MS does not allow direct discrimination between intact NPs and dissolved ions, and the images need to be interpreted according to multi-element localization patterns using tracer elements with a low natural background in plant tissue. For this reason, it remains unclear whether Ce-nPAA-MnO_2_ localized in the proximity of the leaf vasculatures, where they could have started releasing Mn and Ce ions, or whether they were present as intact NPs within them. nPAA-MnO_2_ including a second tracer element (beyond Ce), and possibly phloem immobile, may provide an additional layer of information to help clarifying this. Still, it should be noted that the lateral resolution of LA-ICP-MS (a few microns) may not be sufficient to accurately localize the NPs in the phloem tissue, whose cells are small and tightly packed. Thus, TEM-EDX may be needed to ultimately provide unequivocal proof of intact NPs within the vascular tissues.

### 3.4. Nanoparticle assimilation and mobility

Ionic Ce and Co, with almost-zero ionic background in control plants, were applied to leaves to confirm their contrasting ability to translocate in the phloem (Figure **9**). This experiment confirmed that Ce ions are phloem immobile (Figure 9**B**) whereas Co ions are mobile (Figure **9C**). Four days after the foliar application of Ce/Co labelled nPAA-MnO_2_, both tracer ions largely remained in the exposed leaf (Figures **9B, C**), which shows that the intact NPs was only marginally translocated in the basipetal direction out of the application zone. Comparing the amounts of Ce and Mn in the “Adjacent” fraction upstream of the exposed zone, we could observe that NPs promoted the transport of a small, yet significant, amount of Ce and Mn (0.26% and 1.93%, respectively) away from the exposed area, in comparison to only 0.02% and 0.14% of the applied ionic Ce and Mn. An NP-mediated remobilization of Ce and Mn is consistent with data from the LA-ICP-MS analysis, which highlighted a strong co-localization of Mn and Ce with the leaf vasculature (Figure **8A, B**), suggesting NP presence close to or within the phloem.

The approach utilized here, employing Ce and Co tracers with contrasting phloem mobility, allowed us to obtain information regarding NP intactness, remobilization and dissolution *in planta* (Figure **9**). To the best of our knowledge, this work provides the first example where tracers with different mobility within plants were used to document the assimilation and transport of foliar-applied NPs. Previous studies have reported on the use tracers with low phloem mobility such as Gd^3+^ to study NP translocation in tomato and wheat, but it was never utilized in combination with a mobile tracer like Co.^8,35^ However, owing to different release dynamics of the NP elements *in vitro* settings (Figure **S15**), Ce and Co molar ratios were not preserved over time and thereby it was not possible to model the relative proportions of intact and dissolved NPs. Future studies could address this limitation by employing NP tracer elements and/or stable elemental isotopes with comparable release kinetics and provide deeper insights into NP assimilation and fate within plants.

## 4. Conclusions

Complementary bioimaging and analytical techniques were used to document the uptake and assimilation of foliar-applied nPAA-MnO_2_. Our research highlights the importance of hydraulic activation of the stomatal pathway to enable entry of foliar nPAA-MnO_2_ in barley plants and provides insights into the different transport pathways of mineral ions and nanoparticles across the leaf surface. Following uptake, the pH dependent dissolution of nPAA-MnO_2_ promoted the restoration of Mn deficiency in Mn-deficient barley leaves, showing that foliar nanofertilizers can be used to restore essential metabolic functions. Furthermore, nPAA-MnO_2_ could be applied in high doses without causing leaf scorching, opening new possibilities for increasing the sustainability of foliar fertilization by reducing the frequency of applications and by promoting a gradual release of mineral Mn inside the plant in response to endogenous stimuli, including pH. While showing significant promise as foliar nanofertilizers, this study also highlights major limitations of nPAA-MnO_2_, particularly concerning their ability to remobilize within plants. It was shown that nanotechnology can be used to convert immobile nutrients to mobile ones, but currently to an extent that is irrelevant for agronomic purposes. More research is required to understand the mechanisms controlling phloem loading and long-distance transport of NPs in plants. Understanding these processes represents a major research gap and hampers the design of NP based systems for efficient and systemic delivery of plant nutrients across crop production systems.

## 5. Experimental section

### Plant growth

Spring barley cv. Irina was germinated in vermiculite for 7 days. Uniform seedlings were transferred to aerated, light-impermeable black pots grown under controlled greenhouse conditions.^24^ Each pot (5 L) contained four plants and was filled with a chelate-buffered solution prepared in 18.2 MΩ Milli-Q water (Milli-Q Plus; Millipore). Control plants were grown with sufficient supply of all nutrients, i.e.: KH_2_PO_4_ (200 μM), K_2_SO_4_ (200 μM), MgSO_4_·7H_2_O (300 μM), NaCl (100 μM), Mg(NO_3_)_2_·6H_2_O (300 μM), Ca(NO_3_)_2_·4H_2_O (900 μM), KNO_3_ (600 μM), Fe(III)–EDTA–Na (50 μM), H_3_BO_3_ (2 μM), Na_2_MoO_4_·2H_2_O (0.8 μM), ZnCl_2_ (0.7 μM), MnCl_2_·4H_2_O (1 μM) and CuSO_4_·5H_2_O (0.8μM). The double-ion exchanged water in the buckets was renewed weekly and resupplied with nutrients. The Mn-deficient plants were supplied with a concentration of 10 nM MnCl_2_·4H_2_O on a weekly basis for the first 21 days, which was then increased to 40 nM per unit and supplied every second day. This was adjusted to prevent plant Mn status from declining below the quantum yield threshold for severe Mn deficiency where irreversible tissue damage occurs, defined as Fv/Fm <0.5 measured by chlorophyll *a* fluorescence analysis.^36^

### Characterization of the plant Mn nutritional status

Control and Mn-deficient plant nutritional status was continuously monitored over 31 days by measuring the chlorophyll *a* fluorescence induction kinetics on the YFEL using a Handy PEA chlorophyll fluorometer (Hansatech Instruments), as outlined in Schmidt *et al.*.^37^ Three replicate measurements were made within each independent biological replicate unit. The mid-section of each YFEL was dark-adapted for at least 20 minutes using Hansatech leaf clips before measuring. The fluorescence measurements were recorded by illuminating the leaf with saturating light intensity (3000 μmol photons m^−2^ s^−1^) for 10 s. Plant Mn status was determined by calculating the quantum yield efficiency of PSII as the ratio, Fv(Fm – F0), to F0.^36^

In a separate experiment, the YFEL of control and Mn-deficient barley plants 21 DAT were harvested for Mn tissue analysis. Four independent (n=4) and 3 technical replicates were considered for both the control and the Mn-deficient treatments. Samples were dried at 60 °C for 48 hours, before being digested in a Milestone Ultra WAVE. Samples were analyzed to determine the Mn concentration in the YFEL using ICP-OES (5100 Agilent Technologies).

### Synthesis of nPAA-MnO_2_

nPAA-MnO_2_ were synthesized under N_2_ atmosphere through a one-pot polyol method following concepts of Marasini *et al.,*^38^ with some important modifications. Before the synthesis, in a separate beaker, 10 mmol of NaOH (in tablets) was dissolved in 15 mL of triethylene glycol (TEG) at room temperature for 2 days under constant stirring. For the synthesis, 2 mmol of MnCl_2_·4H_2_O was mixed in 20 mL of TEG in a 100-mL three-neck round-bottom flask equipped with a temperature probe and a glass cold finger. The solution was heated to 80 °C and vigorously stirred for 1 hour to promote the breakdown of MnCl_2_·4H_2_O crystals. Then, the solution was allowed to cool down before 0.25 mmol of PAA (M_w_ = ∼1800 amu; Sigma Aldrich) was added. The solution was then stirred for 4 hours. Afterwards, the 10 mmol NaOH solution was added and the reaction temperature slowly increased from room temperature to 110 °C and maintained for 15 h. Then the solution was cooled to room temperature and transferred to a 500 mL glass media bottle containing 400 mL of 96% ethanol. After 2 days, the sediment containing the NPs was kept, whereas the supernatant was discarded and the ethanol replaced. Three further washing steps with ethanol were performed, replacing the ethanol solution once a week. Finally, NPs were suspended in 100 mL of Milli-Q water and placed into a dialysis tubing (Snakeskin^TM^ Dialysis Tubing 3.5K MWCO; Thermo Fisher Scientific) and dialyzed against 5 L of Milli-Q water, changing the water daily, for 2 days. Ultimately, the NP solution was centrifuged at 40 °C for 8 hours at 25 mbar using a rotary evaporator (Christ Alpha 2-4 RVC) to obtain a volume of 20-30 mL of NP solution.

### nPAA-MnO_2_ characterization

For NP characterization studies, nPAA-MnO_2_ were either: (*i*) kept as an aqueous NP suspension or (*ii*) vacuum-dried to obtain a powder sample.

i. Liquid preparations were used for DLS, Raman spectroscopy, TEM and AFM. The NP hydrodynamic diameter and zeta potential were measured as the average of five consecutive measurements using a Zetasizer Nano ZS (Malvern Panalytical, UK). The hydrodynamic diameter and zeta potential of pristine nPAA-MnO_2_ was measured on five different batches. The Raman spectrum of nPAA-MnO_2_ was acquired using an InVia Raman Spectrometer (Renishaw, UK) with a 532 nm laser source. To record the spectrum, the laser was set to 10 mW power, and the total excitation time was 5s. The sample for the AFM was prepared by putting a drop diluted solution onto a freshly cleaved mica (SPI supplies, grade v-4) surface, letting it stay for 30 seconds and then blow the surface dry with a steady stream of clean nitrogen. For the subsequent imaging, a Cypher from Asylum research (now Oxford instruments) was used. The instrument was equipped with a standard silicon tip, with a spring constant around 2 nN/nm and a resonant frequency around 70 kHz (240AC, Opus *µ*-masch). The image format was set at 512 × 512 pixels. The scanning was performed using AC-mode (tapping) in ambient conditions at 1 Hz. Igor Pro software was used for image acquisition and data analysis. The AFM image was flattened and the thickness (size) of the particles was measured using the section analysis tool in the Igor Pro control software for the AFM. TEM was performed to study NP morphology. First, a carbon-coated Cu-slot grid was prepared using a glow discharger instrument (Leica Coater Ace 200). Then, a 3 µl droplet was deposited on the carbon grid, and the excess liquid was carefully removed using filter paper (Whatman® qualitative filter paper, Grade 1). The imaging was performed using a ThermoFisher Scientific Talos L120C microscope equipped with an energy dispersive spectroscopy (EDS) detector (Bruker XFlash 6T-30) at an accelerating voltage of 120 kV. The EDS spectrum of nPAA-MnO_2_ NPs was recorded in scanning transmission electron microscopy (STEM) mode. The images obtained by AFM and TEM were further processed using FIJI Software (ImageJ) to calculate the particle size distribution.
ii. The dry NP powder was required to perform the TGA and XRD analyses. A thermogravimetric analyzer (Mettler Toledo TGA 2) was used to record changes in NP composition at temperatures ranging from room temperature to 700 °C under an inert N_2_ atmosphere. XRD was performed using an Aeris diffractometer (PANalytical) with Cu k*α* radiation (E = 8.05 keV). Data range was set between 8.005° and 84.998°, with step size 0.011°.

### Preparation of DiI-nPAA-MnO_2_

The procedure for labelling nPAA-MnO_2_ with DiI fluorescent dye (Invitrogen) was adapted from Santra *et al.*.^39^ To a 4 mL nPAA-MnO_2_ suspension (3 mg mL^-1^ Mn), a 500 μL of DiI dye solution (150 μL of DiI, 3 mg mL^-1^, in 350 μL of dimethylsulfoxide (DMSO)) was added dropwise under continuous stirring at ambient temperature. The NPs were kept stirring for 15 minutes. The resulting DiI-nPAA-MnO_2_ were dialyzed against 5 L of Milli-Q water, changing the water daily, for 2 days. To verify the successful encapsulation of DiI dye, the DiI-nPAA-MnO_2_ were analyzed by fluorescence spectrometry using a microplate reader (CLARIOstar Plus, BMG Labtech). The hydrodynamic diameter and zeta potential of the NPs were measured by DLS before and after the fluorescent modification.

### Preparation of Ce-nPAA-MnO_2_ and Co/Ce-nPAA-MnO_2_

Modified nPAA-MnO_2_ including elemental tracers (*i.e.* Ce and Co) were prepared for the NP plant application experiments. Ce-nPAA-MnO_2_ and Ce/Co-nPAA-MnO_2_ NPs were prepared by mixing either 0.02 mmol of Ce(NO_3_)_3_·6H_2_O alone or in combination with 0.02 mmol of CoCl_2_·6H_2_O, together with 2 mmol of MnCl_2_·4H_2_O (molar ratios: Mn:Ce = 1:100; Mn:Ce:Co = 100:1:1, respectively) in the 20 mL TEG mixture at the beginning of the NP synthesis. After the synthesis, NPs were washed in ethanol and then dialyzed in Milli-Q water following the same procedure as for the pristine nPAA-MnO_2_.

To confirm the incorporation of Ce and Co in the final NP product, NPs were dissolved in 3.5% HNO_3_ and analyzed using an 8900 ICP-QQQ-MS (Agilent Technologies).

### Surface tension analysis of the foliar-applied formulations

The surface tension of the foliar-applied solutions formulated with the organosilicone surfactant Silwet Gold, with or without glycerol, was determined by using an optical tensiometer (Theta flow, Biolin Scientific). Silwet Gold surfactant was used to lower the surface tension of foliar formulations, whereas glycerol was used as humectant to improve leaf wetting and extend the droplet drying time of foliar formulations. Briefly, four different solutions containing H_2_O, MnSO_4_, nPAA-MnO_2_ and nPAA-MnO_2_ with 3% glycerol, were prepared with increasing Silwet Gold concentrations (‰ v/v), namely: 0, 0.025, 0.05, 0.075 and 0.1.Thus, a total number of 20 samples were prepared. Three consecutive measurements of 60 second each were conducted for every sample. Values were recorded at every second. Finally, the last five values of each measurement were used for the statistical analysis.

### Dissolution assay

An *in vitro* dissolution assay was set up to study the Mn release kinetics from the NPs at three different physiologically relevant pH values, namely pH 5.5, 6.5 and 7.5. Briefly, 3 mL (1 g L^-1^ Mn) of nPAA-MnO_2_ solution was put inside a dialysis tubing (SnakeSkin™, 3.5K MWCO) and placed in a beaker filled with 1 L of trisodium citrate (Na₃C₆H₅O₇·2H₂O, M_w_= 294,1 g mol^-1^) buffer under stirring for 72 hours. Four technical replicates were considered for each pH value resulting in a total number of 12 beaker. Three mL of the acceptor phase were collected at the time points: 0, 0.5, 1, 2, 4, 8, 24, 48 and 72 hours. At the end of the 72 hours, the volume contained inside each dialysis bag was noted to determine the amount of Mn left. The samples were then diluted in 3.5% HNO_3_ and analyzed using a 5100 ICP-OES (Agilent Technologies). Finally, the mass balance for each sample was calculated and the data were reported as cumulative release (%) over time. The same setup was utilized for the dissolution assay of the double-spiked Ce/Co-nPAA-MnO_2_ NPs in a citrate buffer at pH 5.5.

### Foliar application of common Mn fertilizers and nPAA-MnO_2_

Two different Mn sources and nPAA-MnO_2_ at different concentrations were tested on Mn-deficient leaves to determine their scorching effect. Briefly, four independent treatment stock solutions of MnCl_2_·4H_2_O, MnSO_4_·H_2_O and nPAA-MnO_2_ containing 5 gL^-1^ of Mn were prepared. From these, separate dilutions containing 0.5, 1, 2, 3 and 4 gL^-1^ of Mn were produced. The solutions were then formulated with 0.1‰ Silwet Gold and 3% glycerol. An additional control solution containing only Silwet Gold and glycerol was prepared. For each treatment solution (*i.e.* Mn salts and NPs), 4 μL droplets from every dilution were applied in pairs on the adaxial surface of four replicate YFELs per hydroponic treatment at tillering (21 Days After Transplanting, DAT). In the end, the whole range of Mn concentrations for a given treatment, from 0.5 to 4 gL^-1^, was present on the leaf surface. After 3 days, the leaves were rinsed with Milli-Q water, photographed with a conventional camera and then dark-adapted before being imaged using PAM fluorometry.

### PAM fluorometry analysis

The YFELs that received the foliar treatments were dark-adapted for 30 minutes. Subsequently, leaves were excised and placed inside a Walz IMAGING-PAM M-series MAXI (Heinz Walz GmbH). Leaves were exposed to a saturating light pulse regime to assess the leaf Mn status using the Fv/Fm ratio (Schmidt *et al.*, 2016). Information regarding the spatial distribution of Mn status was obtained using the ImagingWin v2.46i software.

### Effects of formulation properties on the uptake efficiency of foliar solutions

The YFELs of 21 DAT Mn-deficient plants were dosed with four different treatments. For simplicity, these are either listed together with the letters “S+G”, in case they were formulated with both 0.1‰ Silwet Gold and 3% glycerol, or with the “S”, in case they were formulated only with 0.1‰ Silwet Gold. The treatments were: (*i*) water (S+G), (*ii*) 18 mM MnSO_4_ · H_2_O and 0.18 mM Ce(SO_4_)_2_ (S+G), (*iii*) Ce-nPAA-MnO_2_ (S+G), (*iv*) Ce-nPAA-MnO_2_ NPs (S). Mn concentration of 1 g L^-1^ was selected to reflect what is normally applied in conventional agriculture, while minimizing the risk of leaf scorching. Six independent biological replicates were considered for each treatment. 4 μL droplets were applied on the adaxial face of 21 DAT Mn-deficient barley leaves. Three days after the application, droplets were blotted away and leaves were carefully washed with double-distilled water before the harvest. YFELs were dried at 60 °C for 48 hours, before being digested in a Milestone UltraWAVE. Samples were analyzed to determine the Mn and Ce concentrations in the exposed leaves using an 8900 ICP-QQQ-MS (Agilent Technologies). To calculate the uptake efficiency (%), concentration and biomass were used to calculate the amount of element present in the YFEL (m_YFEL_). Then, the following formula was applied:

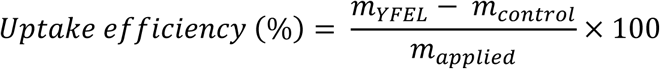

Where *m_control_* and *m_applied_* represent the average amount of element contained in the “Control” treatments (*i.e.* elemental background) and the amount of element applied with the treatment, respectively.

### Confocal imaging of DiI-nPAA-MnO_2_ in plant leaves

To study the leaf uptake of fluorescent DiI-nPAA-MnO_2_, images were collected using a confocal microscope (Leica Stellaris 8). Leaves were treated with DiI-nPAA-MnO_2_ (1 g L^-1^ Mn) formulated with 0.1‰ Silwet Gold and 3% glycerol and then imaged after 2 to 5 hours. Control images were acquired by analyzing barley leaves dosed with an identical formulation containing a 100 times diluted DiI only stock solution (3 mg mL^-1^ in DMSO) instead of DiI-nPAA-MnO_2_. A small portion of the dosed area was excised with a scalpel, placed on a microscope glass slide, and submerged into Perfluorodecalin 95% (Sigma-Aldrich). The white-light laser excitation wavelength was set to 550 nm, and fluorescence was recorded between 560–590 nm and 650–800 nm for DiI and chlorophyll, respectively. Confocal z-stack images were then reconstructed in 3D using the Leica LAS X software.

### Nano-CT imaging of nPAA-MnO_2_ in plant leaves

The nano-CT experiments were carried out at the ID16B beamline of the European Synchrotron Radiation Facility (ESRF) in Grenoble, using the holotomography endstation.^40^ Pristine nPAA-MnO_2_ (1 g L^-1^ Mn) formulate with 0.1‰ Silwet Gold and 3% glycerol were used for the experiments on plants. To test the capabilities of nano-CT, NPs were either (*i*) infiltrated into barley leaves with a syringe, (*ii*) sequentially infiltrated after a 20 mM CaCl_2_ solution or (*iii*) deposited as droplet following infiltration of a 20 mM CaCl_2_ solution. CaCl_2_ was used to induce NP aggregation. A leaf infiltrated with 20 mM CaCl_2_ was included as additional control. 5 hours after treatment, a small rectangular leaf section was placed inside a 200 µL pipette tip and fixed on a rotating sample stage. The beamline was set up in a projection geometry, with a 29 keV pink beam focused to a 50 x 50 nm² spot size using a set of KB mirrors. Each sample was scanned at four different sample-to-detector distances, capturing 900 angles over a 360-degree rotation using a PCO edge detector (2048×2048 pixels). The phase was retrieved using multi-distance Paganin phase retrieval with δ/β = 85 iterative refinement.^41,42^ Data were reconstructed using filtered back projection implemented in PyHST2,^43^ followed by post-processing in Dragonfly, which included 3D shading compensation with the manual radial basis correction function and denoising with a median filter. NP aggregates were segmented for 3D visualization through thresholding and rendered in Dragonfly together with the leaf tissue, while 2D slices were visualized using FiJi, an open source platform for biological image analysis.^44^

### LA-ICP-MS analysis of leaves exposed to Ce-nPAA-MnO_2_

LA-ICP-MS was used to study the leaf distribution of Ce-nPAA-MnO_2_ NPs in the early hours following application. Ce-nPAA-MnO_2_ (1 g L^-1^ Mn) formulated with 0.1‰ Silwet Gold and 3% glycerol were applied to the YFEL of 21 DAT Mn-deficient plants. After 5 hours, the NP droplets were gently removed using a wet paper towel, and then a small leaf area was excised with a scalpel, embedded in OCT and frozen in dry ice-cooled liquid hexane. The OCT molds were sliced at -30°C using a cryotome (Leica CM3050S) and freeze-dried overnight. Thin 14 µm cross sections were ablated using a nanosecond LA unit (Iridia 193 nm excimer laser ablation system, Teledyne CETAC technologies) with the following settings: fluence: 1.3 J cm^-2^; scan speed: 240 μm s^-1^; repetition rate: 300 Hz; and spot size: 4 or 5 μm. An 8900 ICP-QQQ-MS (Agilent Technologies) was used to obtain the element signals, operating in standard mode. The monitored isotopes were ^24^Mg, ^55^Mn and ^140^Ce (integration times for 4 and 5 µm spot size, respectively: 0.4 ms and 0.5 ms for Mg,, 7.8 and 8 ms for Mn, 8.8 ms and 8.5 ms for Ce, respectively). The ICP-MS was operated with sample cone depth = 3 mm and carrier gas = 0.73 L min^-1^. Image processing was performed in HDIP (High-definition Image processing, Teledyne CETC technologies), including gas blank.

### Assimilation and distribution of Ce/Co-nPAA-MnO_2_

A NP leaf application study was designed to assess NP dissolution and translocation dynamics inside the plant. Smaller Mn-deficient barley plants were used for this experiment in order to increase the relative proportion of applied Mn *vs* the plant total Mn (*i.e.* the plant background), thereby facilitating the potential detection of small Mn amounts translocated to unexposed plant parts by ICP-MS. The normal plant growth protocol was used with few modifications. Briefly, barley plants were germinated for 5 days in vermiculite (instead of 7) and then were transplanted into hydroponics pots, where they were grown for 14 days (instead of 21) without Mn supply prior to the experiment.

For the application experiment, three treatments were considered: (*i*) formulation only, (*ii*) ionic solution containing 18 mM MnSO_4_ · H_2_O, 0.18 mM Ce_2_(SO_4_)_3_·8 H_2_O and 0.18 mM CoCl_2_·6 H_2_O and (*iii*) Ce/Co-nPAA-MnO_2_ NPs. NPs were applied at 1 g L^-1^ (= 18mM) Mn concentration. The ionic solution was prepared accordingly, to equal the Mn, Co and Ce concentrations present in the NP solution. All the final solutions were formulated with 0.1‰ Silwet Gold and 3% glycerol. A single YFEL from each plant was dosed, and six independent biological replicates were considered for each treatment. A total of 48 µL (12 droplet of 4 µL) were applied on the adaxial face of 21 DAT Mn-deficient barley leaves. Plants were harvested four days after NP leaf application and were divided into four different fractions: (*i*) exposed area, (*ii*) adjacent area, (*iii*) shoots, and (*iv*) roots. Plant parts were dried at 60 °C for 48 hours, before being homogenized and digested in a Milestone UltraWAVE. Samples were analyzed to determine the concentration of Mn, Ce and Co in the four different fractions using an 8900 ICP-QQQ-MS (Agilent Technologies, USA).

The amount of element translocated to the “Adjacent” fraction was calculated as a percentage of the aggregate amount recovered in the “Exposed” and “Adjacent” fractions, together representing the whole YFEL, using this formula:

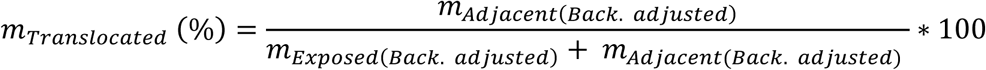

*m_Adjacent(Back. adjusted)_* represents the amount (µg) of element in the “Adjacent” fraction, and it was calculated from the concentration detected and the biomass, and then subtracted by the average elemental background (*i.e.* the “Control” treatment). *m_Exposed(Back. adjusted)_* instead represents the amount (µg) of element in the “Exposed” fraction, and it was calculated from the concentration detected and the biomass, and then subtracted by the average elemental background (*i.e.* the “Control” treatment).

### Statistical analysis

All data were represented as mean, ± indicates SD or SE, and *n* = independent biological replicates. Data visualization and statistical analysis were performed using Rstudio software. Statistical comparisons were performed by unpaired *t-test* or one-way ANOVA based on post-hoc Tukey test using lme4, nlme and emmeans packages. Statistical significances among experimental groups were denoted by different letters or asterisks. All models were validated by evaluation of predicted residuals compared to measured datapoints and by ggplot to ensure normal distribution.

## Acknowledgements

This work was financially supported by the Novo Nordisk Foundation, NNF Challenge Program: Smart Nanomaterials for Applications in Life-Science (Grant: NNF21OC0066114). The European Synchrotron Radiation Facility (ESRF) is acknowledged for provision of beam time (experiment LS3284) using the ID16B beamline.

## Author contributions

AP and SH designed the research. AP, AKG and AS performed the experiments with analytical contributions from BM, JCG (XPS, XRD), NT (LA-ICP-MS), TH (AFM), EVK and RM (nano-CT). AP, SH analyzed data and drafted the paper. All authors contributed to finalizing the paper and approved it before submission.

## Associated content

### Supporting information available

Mn concentration in 21 DAT barley YFEL, AFM image of nPAA-MnO_2_, hydrodynamic diameter and zeta potential of pristine nPAA-MnO_2_ and different modifications, TGA analysis of nPAA-MnO_2_, XRD pattern of nPAA-MnO_2_, TEM and STEM analysis of nPAA-MnO_2_, Raman spectrum of nPAA-MnO_2_, TEM images of Ce-nPAA-MnO_2_ and Ce/Co-nPAA-MnO_2_, fluorescence emission spectra and DLS analyses of DiI-nPAA-MnO_2_, dissolution profiles of nPAA-MnO_2_, surface tension of foliar formulations, uptake efficiency of foliar-applied ions versus Ce-nPAA-MnO_2_, droplet drying time of foliar Ce-nPAA-MnO_2_ formulations, CLSM videos and control images, nano-CT videos, elemental maps of Mn-deficient barley leaf cross sections, dissolution profiles of Co/Ce-nPAA-MnO_2_, Mn and tracers partitioning in different plant fractions after foliar Co/Ce-nPAA-MnO_2_ exposure.

