## Supplemental Information for "Foliar Application of Polymer-coated Manganese Dioxide Nanoparticles: Mechanisms of Uptake and Metabolic Responses in Manganese Deficient Barley"

### Supporting Information

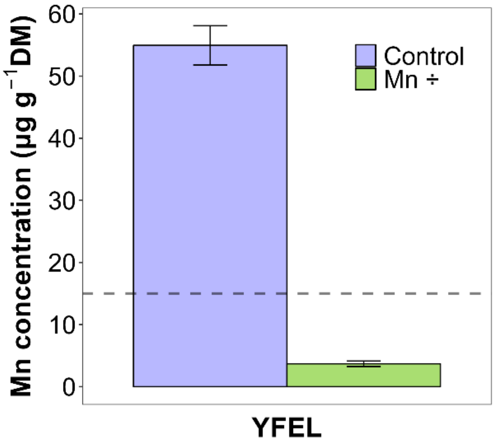


**Figure S1**: Mn concentration in 21 DAT barley YFEL. The purple bar shows control leaves, whereas the green bar shows the Mn-deficient leaves. The dotted line, set at 15 µg $g^{-1}$ DM, indicates the threshold for Mn-deficiency. Results are presented as means ± SD (n=4 with 3 technical replicates).

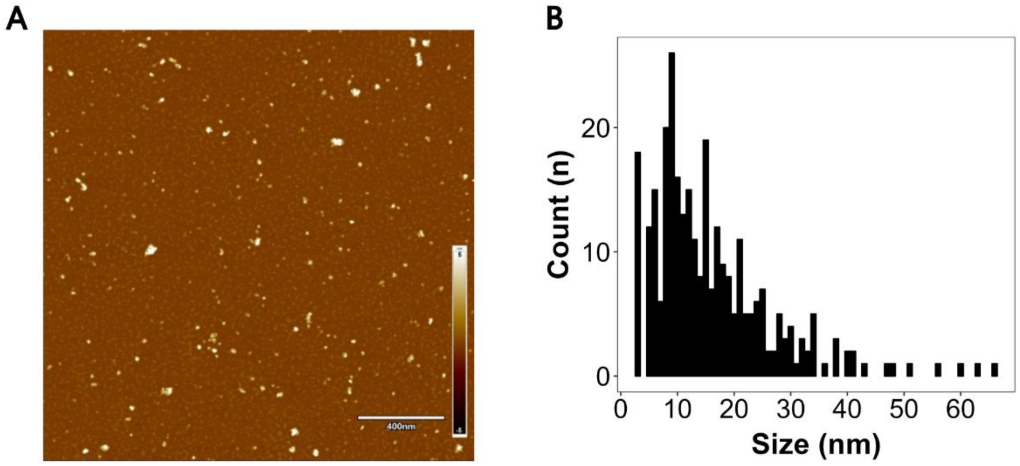


**Figure S2**: (A) 2x2 µm AFM topography image of a nPAA-$MnO_2$ sample on a freshly cleaved mica support. Small particles ranging from about 3 to 15 nm in diameter are consistent with nPAA-$MnO_2$ single native particles observed in TEM (Figures **2B**, **C**). Bigger particles, up to 65 nm and irregularly shaped, are consistent with nPAA-$MnO_2$ aggregates. Small particles with sub-nm size (0.3-0.6 nm) are visible on the background and are consistent with patches of PAA molecules likely resulting from the drying of the NP sample. (B) Size distribution plot obtained from the AFM image shows the highest frequency count at 9 nm, in consistency with the TEM data (n=321).

**Table S1**: Hydrodynamic diameter and zeta potential of pristine nPAA-$MnO_2$ and different modifications, including Ce-nPAA-$MnO_2$ and Ce/Co-nPAA-$MnO_2$, measured by DLS.

| Sample | Size (nm) | Zeta (mV) |
|---|---|---|
| Pristine nPAA-$MnO_2$ | 25.9 ± 5.3* | -46.4 ± 3.7* |
| Ce-nPAA-$MnO_2$ | 18.97 ± 0.23 | -50.7 ± 1.1 |
| Ce/Co-nPAA-$MnO_2$ | 20.78 ± 0.14 | -40.5 ± 1.1 |

* The pristine nPAA-$MnO_2$ was measured across five distinct batches. In the cases of Ce-nPAA-$MnO_2$ and Ce/Co-nPAA-$MnO_2$, the results reflect the average of five sequential measurements taken from a single sample, as only one batch was synthesized for each.

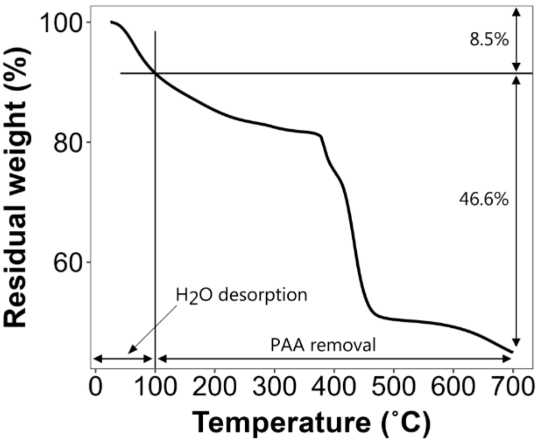


**Figure S3**: Thermogravimetric analysis of nPAA-$MnO_2$ performed at a temperature range between 25°C and 700°C. The initial weight loss (approximately up to 350 °C) can be attributed to dehydration and decarboxylation of PAA. The further weight loss reduction can be attributed to the complete degradation of PAA oligomers.

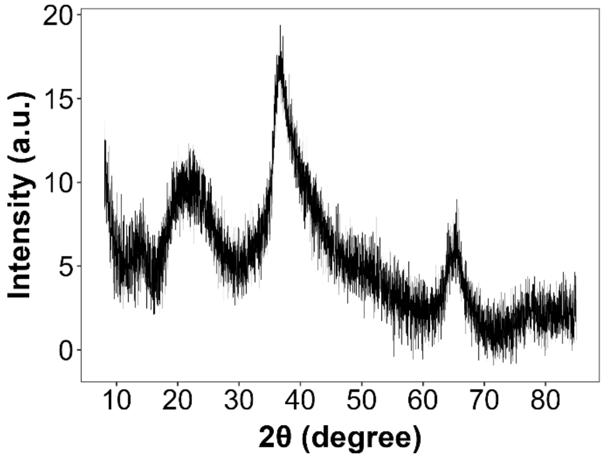


**Figure S4**: Experimental XRD pattern of a powdered nPAA-$MnO_2$ sample. A broad hump extending between ~50 and 55 is characteristic for layered $\delta$-$MnO_2$ (birnessite) phase.[14]

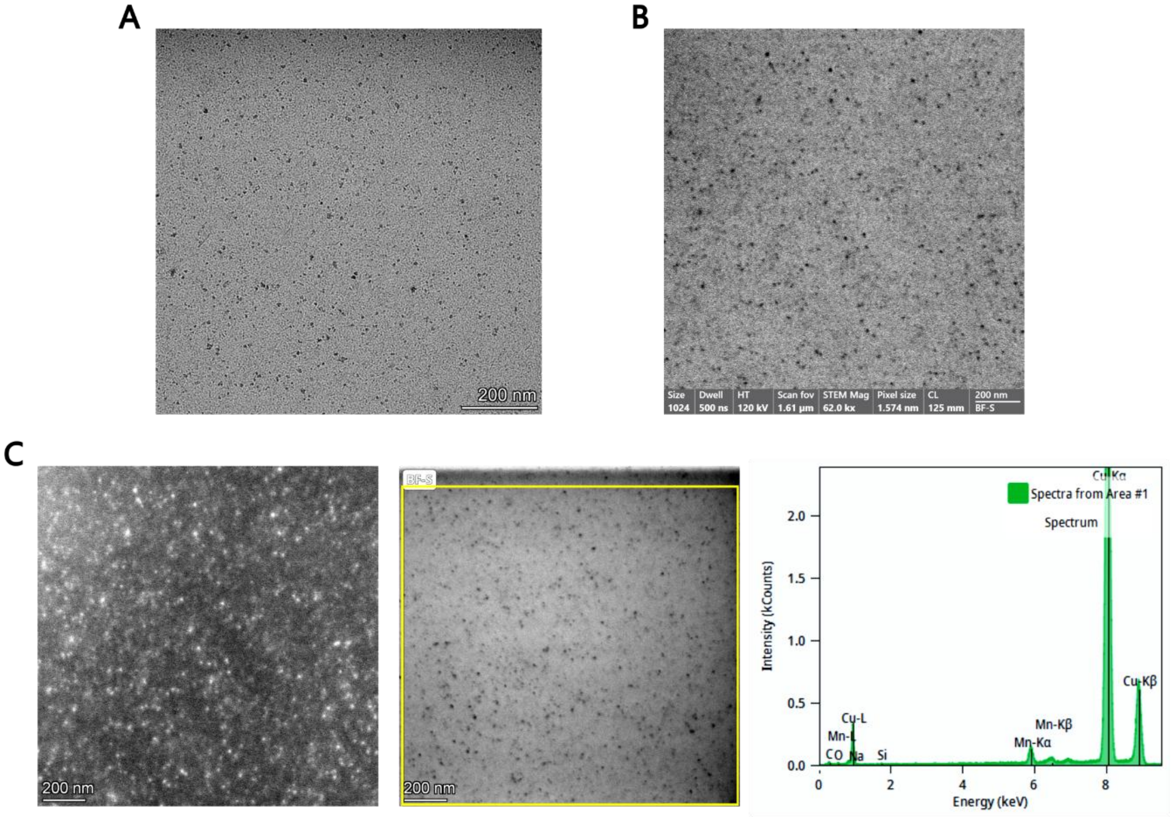


**Figure S5**: (A) TEM image nPAA-$MnO_2$. (B) STEM image of nPAA-$MnO_2$. (C) STEM analysis of nPAA-$MnO_2$. From *left* to *right*: dark-field image showing the back-scattered electron of nPAA-$MnO_2$, corresponding bright-field image and EDS spectrum collected in the area delimited by the yellow rectangle.

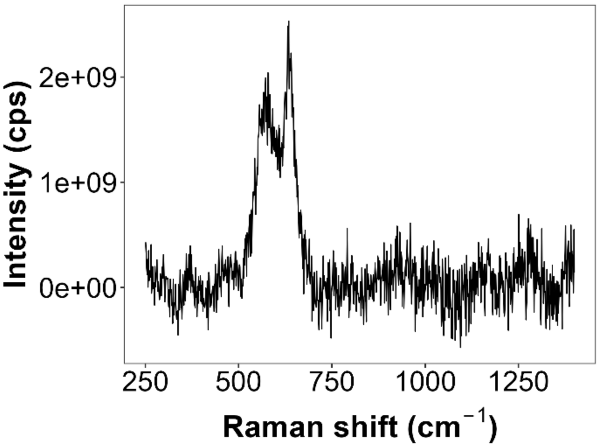


**Figure S6**: Raman spectrum of a liquid nPAA-$MnO_2$ sample. Characteristic peaks 637 $cm^{-1}$ and 580 $cm^{-1}$ can be attributed to to the Mn–O stretching vibration of the $MnO_6$ octahedra and the basal plane of the $MnO_6$ sheet of birnessite, respectively.[16,17]

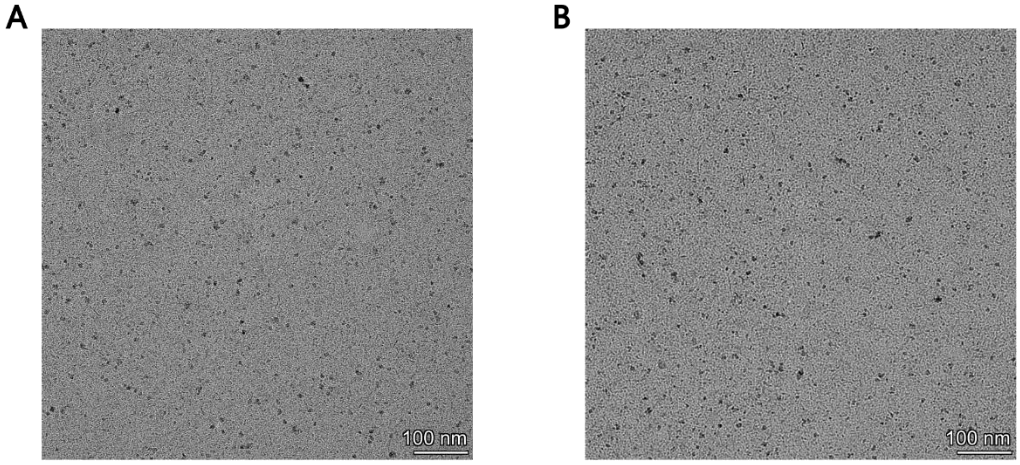


**Figure S7**: Ce-nPAA-$MnO_2$ (A) and Ce/Co-nPAA-$MnO_2$ (B) TEM images show similar size and morphology of the pristine nPAA-$MnO_2$. Due to their presence in small quantities (Ce:Co:Mn molar ratio is ~1:1:100), Ce and Co could not be detected in EDS.

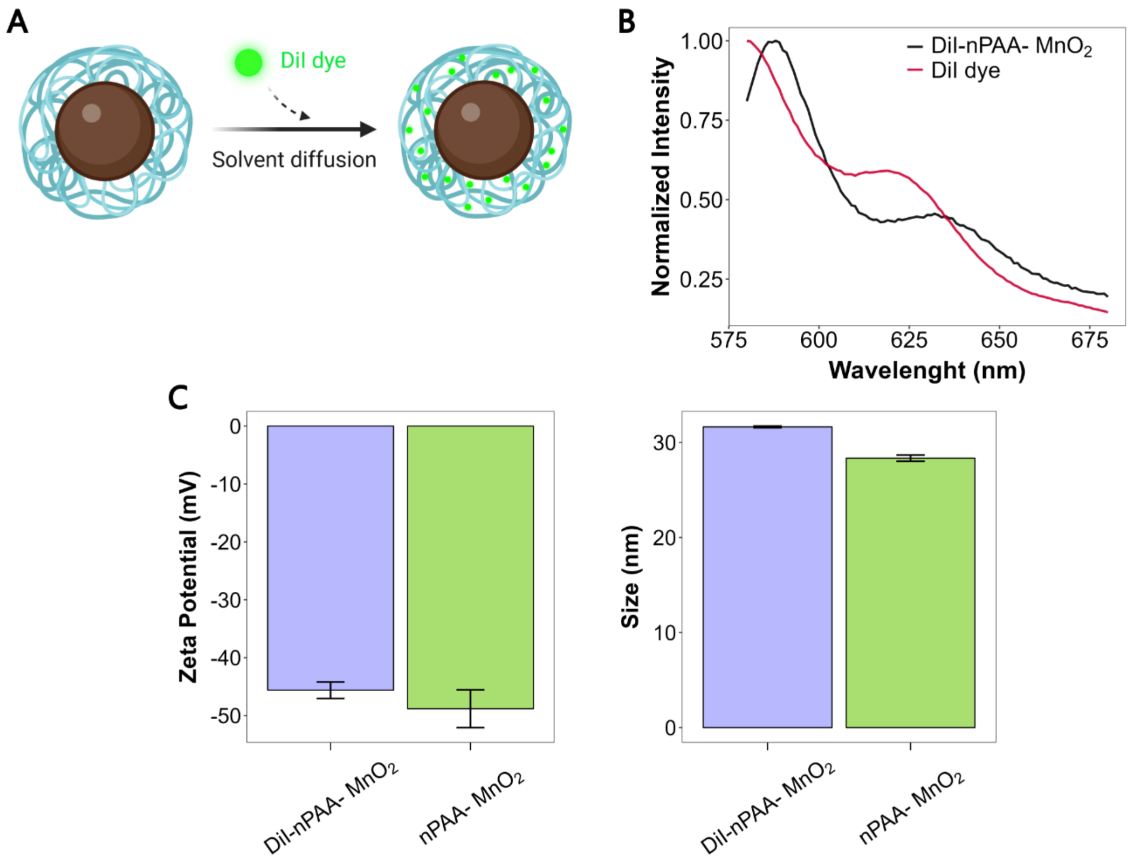


**Figure S8**: (A) Schematic representation of dye-encapsulation method utilized to label nPAA-$MnO_2$. This approach does not alter the stability of NPs in aqueous media, nor their surface properties as the dye is stably incorporated into the polymeric coating. (B) Fluorescence emission spectra of the DiI-nPAA-$MnO_2$ and DiI dye alone. The encapsulation of DiI in the NP coating was confirmed by the presence of a red-shift in the fluorescence peak of the DiI-nPAA-$MnO_2$ compared to the free DiI.[39] (C) Zeta potential and hydrodynamic size of nPAA-$MnO_2$ before and after labelling with the DiI dye measured by DLS. Five consecutive measurements were performed on each sample.

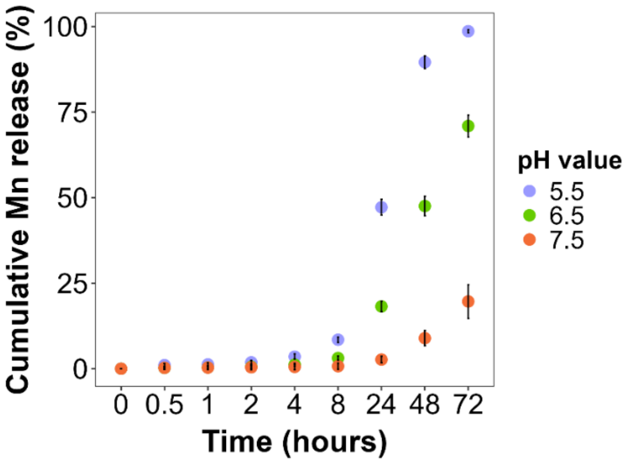


**Figure S9**: Dissolution profiles of nPAA-$MnO_2$ in 5 mM citrate buffer at pH 5.5, 6.5 and 7.5. Results are presented as means ± SD (n=1 with 4 technical replicates).

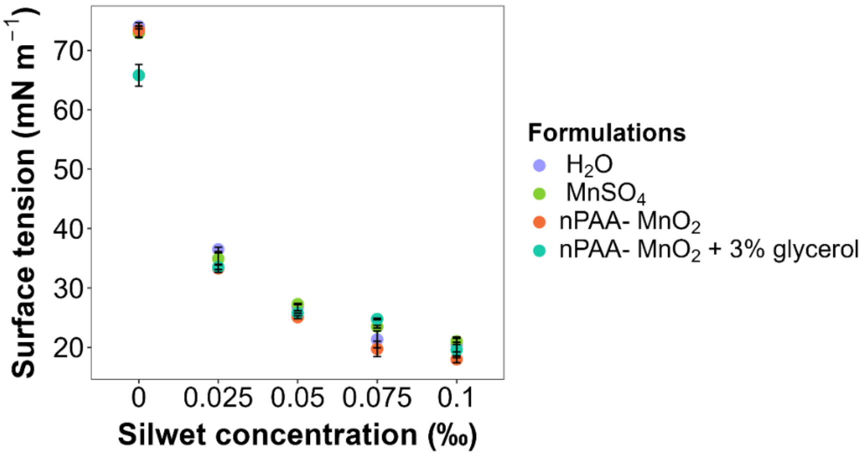


**Figure S10**: Surface tension of different foliar solutions at increasing concentration (‰) of Silwet gold surfactant.

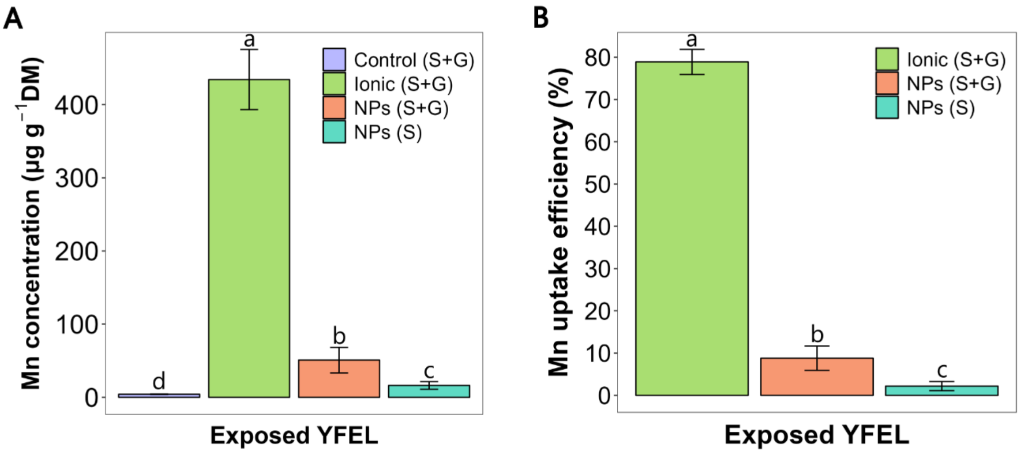


**Figure S11**: Uptake efficiency of foliar-applied ions versus Ce-nPAA-$MnO_2$, formulated with or without 3% glycerol. YFELs were sampled 3 days after NP exposure. (A) Mn concentration in the exposed YFEL. (B) Uptake efficiency of Mn in the different treatments measured as a percentage of the total amount applied. Results are presented as means ± standard deviation of the mean ($n = 6$), and letters represent significant differences ($p < 0.05$) analyzed by a one-way ANOVA and Tukey's multiple comparison test.

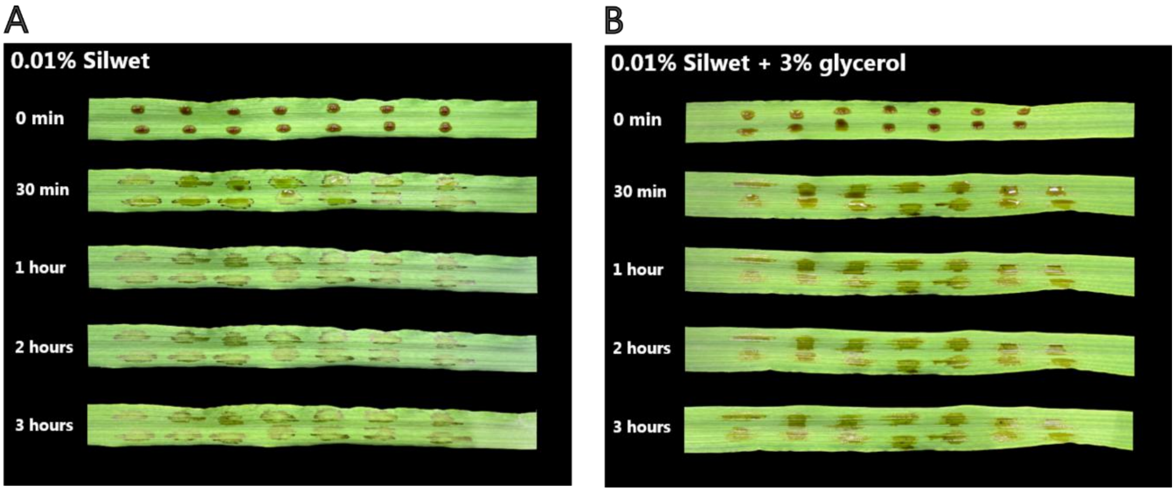


**Figure S12**: Droplet drying time. Time sequence of foliar-applied Ce-nPAA-$MnO_2$ formulated with either Silwet Gold only (A) or in combination with glycerol (B).

CLSM videos are available here

**Video S1:** 3D visualization of DiI-nPAA-$MnO_2$ uptake 2 hours after leaf application. In red: chlorophyll autofluorescence from the chloroplasts; in green: DiI signal.

**Video S2:** 3D visualization of foliar-applied DiI dye formulated with 0.1‰ Silwet and 3% glycerol. In red: chlorophyll autofluorescence from the chloroplasts; in green: DiI signal.

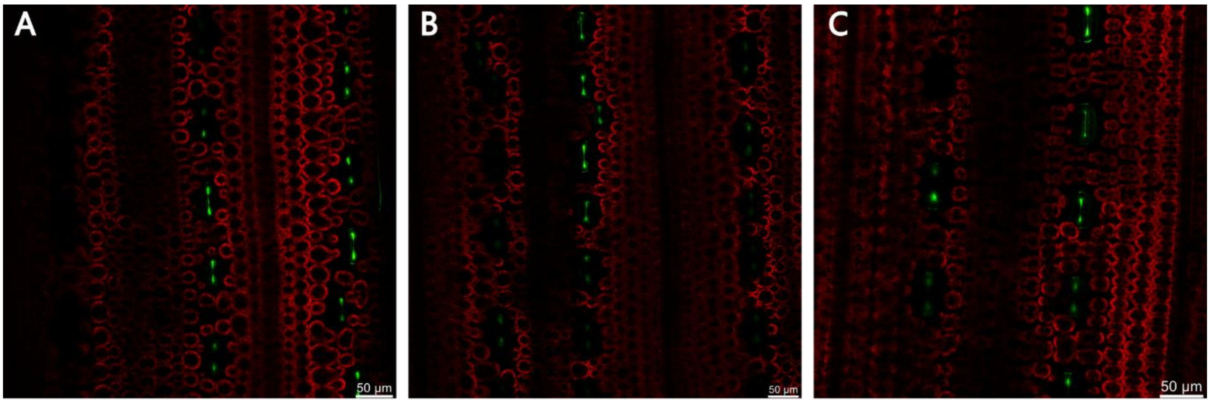


**Figure S13**: Confocal images of barley leaf treated with a DiI dye formulation containing 0.1‰ Silwet Gold and 3% glycerol. Images (A), (B) and (C) were taken 3, 4 and 5 hours after the leaf application, respectively. In red: chlorophyll autofluorescence from the chloroplasts; in green: DiI signal. No DiI signal could be detected inside the substomatal cavity.

Nano-CT videos are available here

**Video S3:** 3D visualization of a barley leaf 5 hours after infiltration of nPAA-$MnO_2$ (low resolution, pixel size = 257 nm).

**Video S4:** 3D visualization of a barley leaf 5 hours after infiltration of nPAA-$MnO_2$ (high resolution, pixel size = 50 nm).

**Video S5**: 3D visualization of a barley leaf where nPAA-$MnO_2$ were sequentially infiltrated after 20 mM $CaCl_2$ (high resolution, pixel size = 50 nm). NP clusters are visible in red.

**Video S6**: 3D visualization of nPAA-$MnO_2$ sequentially infiltrated after 20 mM $CaCl_2$ near the stomata region (high resolution, pixel size = 50 nm). NP clusters are visible in red.

**Video S7**: 3D visualization of a barley leaf 5 hours after $CaCl_2$ infiltration and nPAA-$MnO_2$ deposition on the leaf surface (low resolution, pixel size = 257 nm). NP clusters are visible in red.

**Video S8**: 3D visualization of a barley leaf infiltrated with 20 mM $CaCl_2$ (low resolution, pixel size = 257 nm).

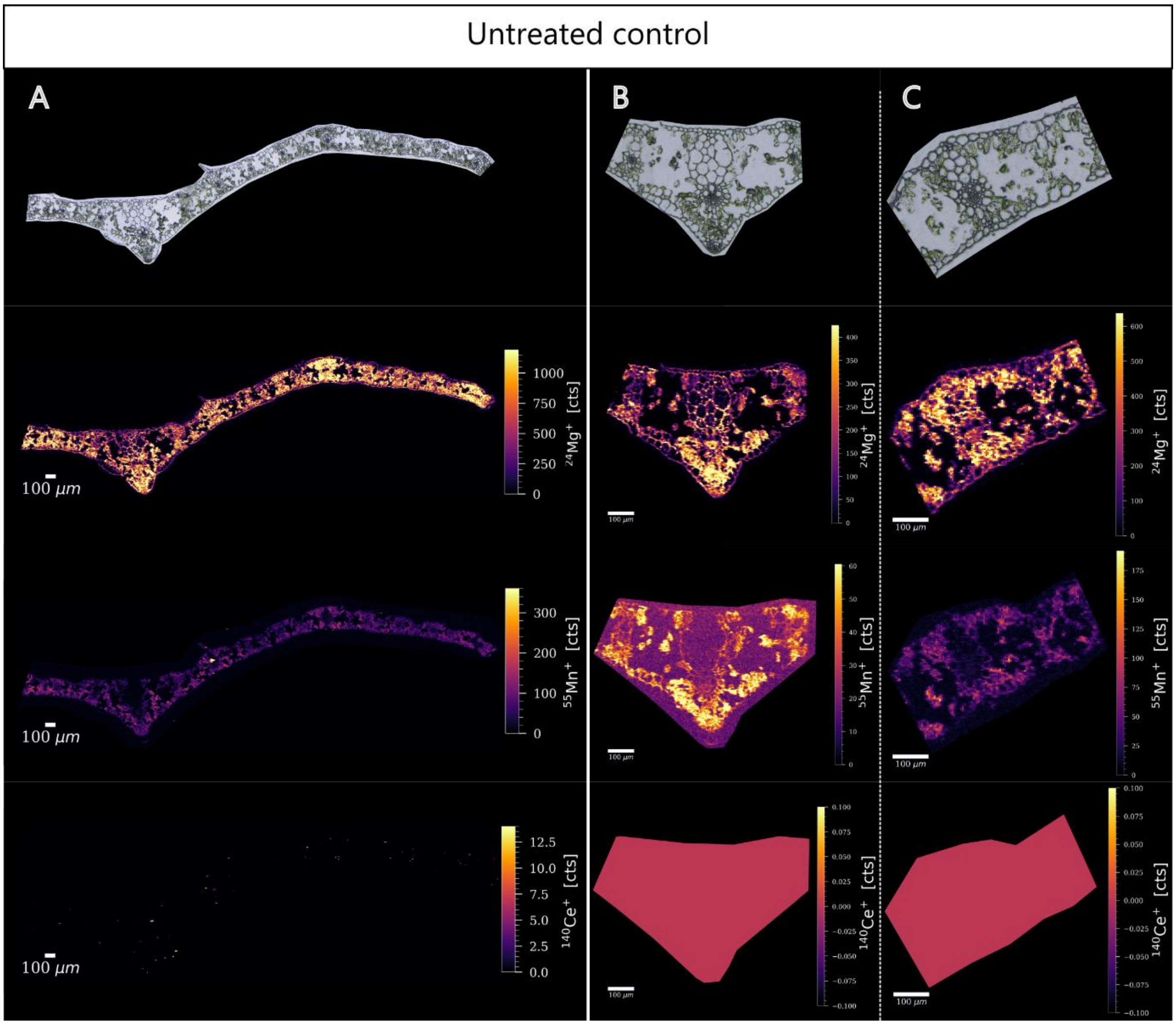


**Figure S14**: Elemental maps of 14 µm-thick Mn-deficient barley leaf cross sections. (A) Leaf overview (spotsize 5 µm). (B) and (C) close-ups of leaf vasculature (spotsize 4 µm). Mg is ubiquitously present in plant tissues and it was used to highlight the plant CWs. Mn and Ce where mapped to assess the background in plants.

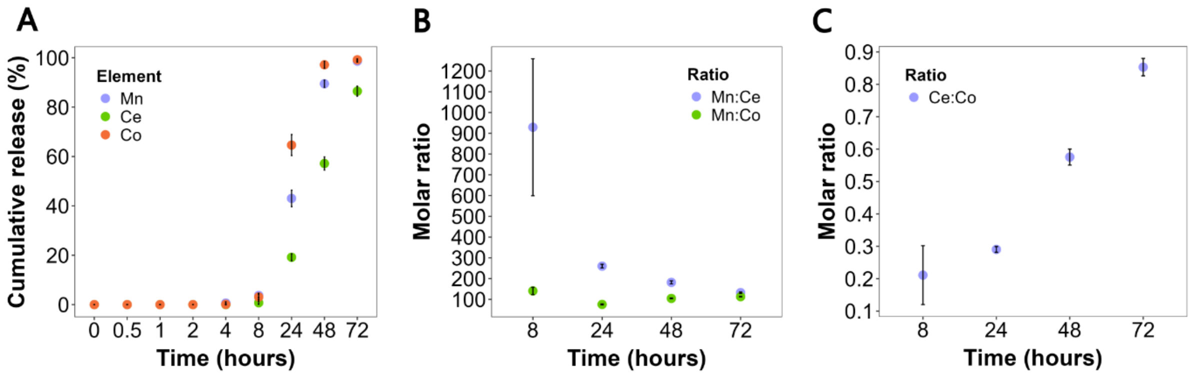


**Figure S15**: dissolution profiles of Co/Ce-nPAA-$MnO_2$ in 5 mM citrate at pH 5.5. (A) Cumulative element release of Mn and Ce, Co tracers. (B) Molar ratio profiles of Mn:Ce and Mn:Co. (C) Molar ratio profile of Ce:Co tracers.

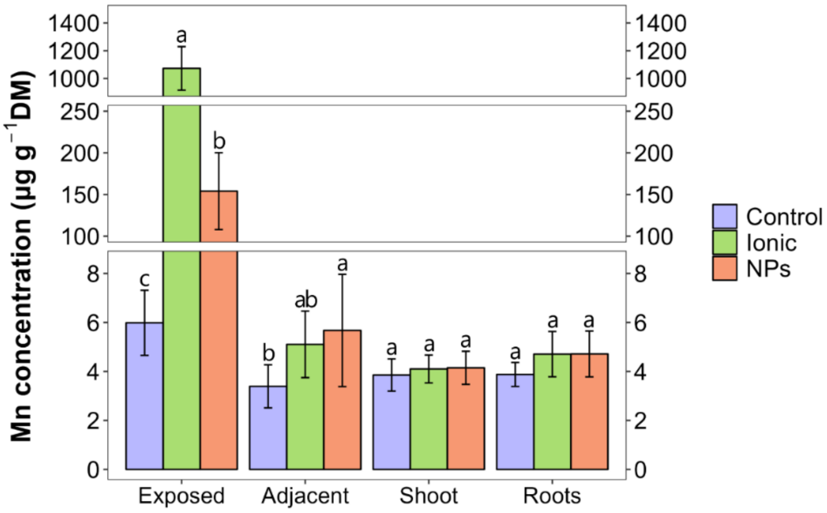


**Figure S16**: Mn distribution in different plant fractions 4 days after foliar exposure to Ce/Co-nPAA-$MnO_2$ and an analogous ionic solution. The legend shows the three different treatment consisting of different formulations, namely: Silwet Gold and glycerol (*Control*), $MnSO_4$, $CeSO_4$ and $CoCl_2$ (*Ionic*), and Ce/Co-nPAA-$MnO_2$ (*NPs*). Results are presented as means ± standard deviation of the mean ($n = 6$), and letters represent significant differences ($p < 0.05$) analyzed by a one-way ANOVA and Tukey's multiple comparison test carried out within groups.
